# Unlocking Antarctic molecular time-capsules – recovering historical environmental DNA from museum-preserved sponges

**DOI:** 10.1101/2024.04.28.591519

**Authors:** Gert-Jan Jeunen, Sadie Mills, Miles Lamare, Grant A. Duffy, Michael Knapp, Jo-Ann L. Stanton, Stefano Mariani, Jackson Treece, Sara Ferreira, Benjamín Durán-Vinet, Monika Zavodna, Neil J. Gemmell

**Affiliations:** Department of Marine Science, University of Otago, Dunedin, New Zealand; Department of Anatomy, University of Otago, Dunedin, New Zealand; National Institute of Water & Atmospheric Research, Wellington, New Zealand; Liverpool John Moores University, Liverpool, UK; Otago Genomics Facility, University of Otago, Dunedin, New Zealand

**Keywords:** metabarcoding, fish diversity, Porifera, ethanol DNA extraction, frozen DNA extraction, dried DNA extraction

## Abstract

Marine sponges have recently emerged as efficient natural environmental DNA (eDNA) samplers. The ability of sponges to accumulate eDNA provides an exciting opportunity to reconstruct contemporary communities and ecosystems with high temporal and spatial precision. However, the use of historical eDNA (heDNA), trapped within the vast number of specimens stored in scientific collections, opens up the opportunity to begin to reconstruct the communities and ecosystems of the past. Here, using a variety of Antarctic sponge specimens stored in an extensive marine invertebrate collection, we were able to recover information on Antarctic fish biodiversity from specimens up to 20 years old. We successfully recovered 64 fish heDNA signals from 27 sponge specimens. Alpha diversity measures did not differ among preservation methods, but sponges stored frozen had a significantly different fish community composition compared to those stored dry or in ethanol. Our results show that we were consistently and reliably able to extract the heDNA trapped within marine sponge specimens, thereby enabling the reconstruction and investigation of communities and ecosystems of the recent past with a spatial and temporal resolution previously unattainable. Future research into heDNA extraction from other preservation methods, as well as the impact of specimen age and collection method will strengthen and expand the opportunities for this novel resource to access new knowledge on ecological change during the last century.

## 1 INTRODUCTION

Environmental DNA (eDNA) surveys have revolutionised how scientists monitor the Earth’s marine biome (Takahashi et al. 2023). The capacity to discern biodiversity and ecological processes utilising genetic material extracted from environmental samples, such as water (Cecchetto et al. 2021), soil (Olmedo-Rojas et al. 2023), sediment (Kuwae et al. 2020), air (Lynggaard et al. 2022), or gut content (Vasiliadis et al. 2024), obviates the necessity for direct species observations, a challenging accomplishment for the inaccessible and vast marine environment (Takahashi et al. 2023). Hence, eDNA metabarcoding surveys detect a significantly larger proportion of the marine biological community compared to traditional approaches, such as diver surveys (Robinson et al. 2023), baited remote underwater video (Stat et al. 2019), and trawling (Maiello et al. 2022). While a partial overlap in species detection is most commonly observed in comparative experiments with traditional monitoring approaches (Robinson et al. 2023), eDNA species detection reliability will further increase when overcoming current limitations, such as enhanced primer design (Wang et al. 2023) and more complete reference databases (Stoeckle et al. 2020). The application of eDNA metabarcoding, therefore, has the potential to increase species detection efficiency while also offering the advantage of non-invasive sampling to reduce potential disturbances to fragile marine ecosystems (Takahashi et al. 2023).

Aquatic eDNA surveys have been observed to achieve high spatial and temporal resolutions (Minamoto et al. 2017; O’Donnell et al. 2017; Jeunen et al. 2019a; Jensen et al. 2022), thereby enabling accurate species detection of organisms present near the sampled area. This resolution has been linked to high degradation rates of DNA in the environment and influenced by biotic, e.g., bacterial activity (Tsuji et al. 2017), and abiotic factors including pH and temperature (Strickler et al. 2015; Tsuji et al. 2017). The rapid degradation of eDNA in the open marine environment, however, also limits aquatic eDNA surveys to monitoring contemporary biodiversity patterns (Ramírez-Amaro et al. 2022).

Effective conservation of the marine biome requires current biodiversity trends to be interpreted against accurate historical ecological baselines, allowing an understanding of the magnitude and drivers of past changes (Lotze and Worm 2009; Harnik et al. 2012; Finnegan et al. 2015). In terrestrial systems, a wealth of historical data has refined our understanding of the changes brought about by direct (Roberts et al. 2017; Wood et al. 2017) and indirect (Rick et al. 2013; Parducci et al. 2019) human pressures. Marine conservation efforts, on the other hand, have only recently begun to use various historical and ancient data sources to determine ecological baselines for the marine environment, such as fossils (Finnegan et al. 2015), midden remains (Seersholm et al. 2018), sediment cores (Finney et al. 2002), and written records (Pauly and Zeller 2016). Such data sources for the marine biome, however, are extremely scarce (Willis et al. 2007), as well as difficult and expensive to obtain (Kittinger et al. 2015). Furthermore, information on how marine environments have responded to anthropogenic pressures is mostly incomplete (Hoegh-Guldberg and Bruno 2010; Norris et al. 2013; Kidwell 2015). The lack of accurate historical ecological baseline information is particularly pronounced for polar regions, which have suffered profound anthropogenic impacts during the last century through fishing (Pinkerton and Bradford-Grieve 2014), whaling (Aronson et al. 2011), and climate change (Parkinson 2019).

Recently, filter-feeding organisms have been investigated as natural eDNA samplers (Mariani et al. 2019; Junk et al. 2023). In particular, marine sponges have been shown to naturally accumulate environmental DNA by continuously filtering large volumes of water to capture particulate matter as a food source (Godefroy et al. 2019). Compared to aquatic eDNA, marine sponges have been observed to hold near-identical vertebrate and eukaryotic diversity patterns within small spatial scales (Jeunen et al. 2023a, 2023b), as well as mirroring temporal resolutions in a controlled mesocosm experiment (Cai et al. 2022). Similarly to comparisons between aquatic eDNA and traditional survey approaches, a partial overlap between sponge eDNA and visual surveys has been observed, with sponge eDNA recovering a larger fraction of the fish community in deep-sea and polar regions (Brodnicke et al. 2023; Jeunen et al. 2024). The observed variability in the efficiency of capturing and retaining eDNA signals across species within the phylum Porifera (Cai et al. 2022; Brodnicke et al. 2023) has been linked to microbial activity (Brodnicke et al. 2023). The ability of marine sponges to accumulate eDNA through their filter-feeding strategy enables an exciting opportunity to reconstruct past ecosystems with a previously unattainable temporal and spatial precision by extracting historical eDNA (heDNA) from museum-stored sponge specimens (Neave et al. 2023).

While vast numbers of marine sponges have been gathered over centuries for research purposes, various preservation methods have been employed to archive specimens in scientific collections (Ghiglione et al. 2018). For example, within the NIWA Invertebrate Collection (NIC) in New Zealand, marine sponge specimens are most often stored in ethanol, dried, or frozen (FIG 1). A wealth of molecular research aimed at extracting host DNA from museum specimens has revealed preservation techniques to influence DNA degradation rates (Martínková and Searle 2006; Iyavoo et al. 2019), DNA integrity (Zimmermann et al. 2008; Moreau et al. 2013), and laboratory protocol choice (Nishiguchi et al. 2002; Nagy 2010; Rowe et al. 2011; Hahn et al. 2021). Hence, to enable heDNA signal comparisons to be made from sponge specimens stored using different preservation techniques, it is essential to understand how preservation method choice impacts and potentially biases heDNA recovery success.

**FIG 1:**
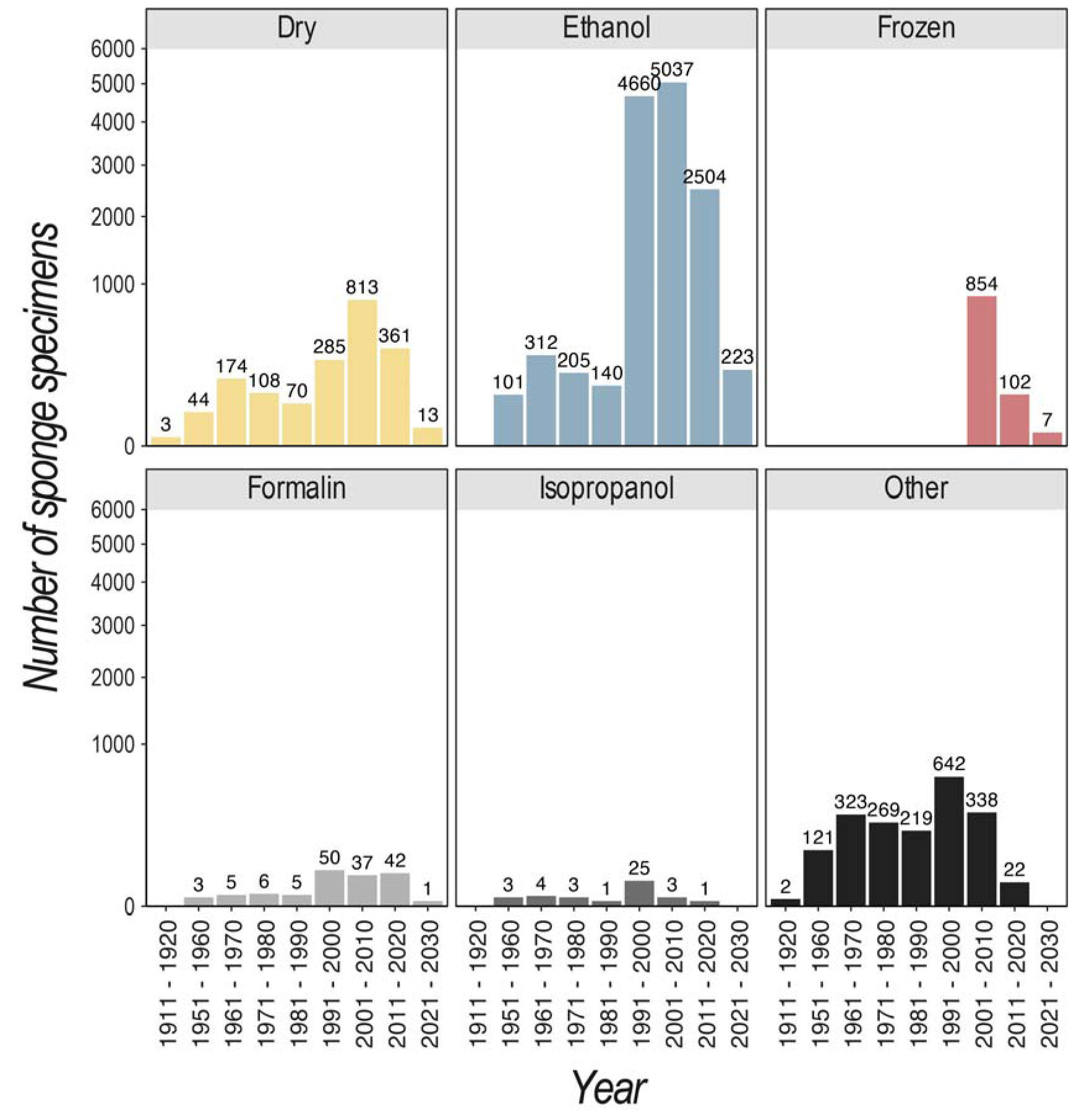
The number of sponge specimens in the NIWA Invertebrate Collection (NIC) per decade and facetted by preservation method, including dry (yellow), ethanol (blue), frozen (red), formalin (light-grey), isopropanol (grey), and other (dark-grey). Specimens included in ‘other’ include preservation methods listed as Alcohol Unknown, Ethanol – Previously Unknown, and Slide. Number above bars represent number of specimens. Y-axis reported as square root transformed to increase readability of low-abundant collection numbers. For NIC specimen data, see https://nzobisipt.niwa.co.nz/resource?r=obisspecify.

In this study, we determine the feasibility of extracting historical fish eDNA signals from 30 Antarctic sponge specimens stored either by ethanol submersion, dried, or frozen (FIG 2). Sponge specimen collection dates ranged from 1960 to 2011. Additionally, we explore the potential bias that preservation methods might introduce to heDNA recovery by comparing alpha and beta diversity metrics from the 30 sponge specimens, while accounting for specimen age and sponge taxonomic ID as potential covariates. Finally, we estimate the replication required to detect 90% of historical fish eDNA signals based on inter- and extrapolation calculations within five tissue biopsy replicates within each sponge specimen.

**FIG 2:**
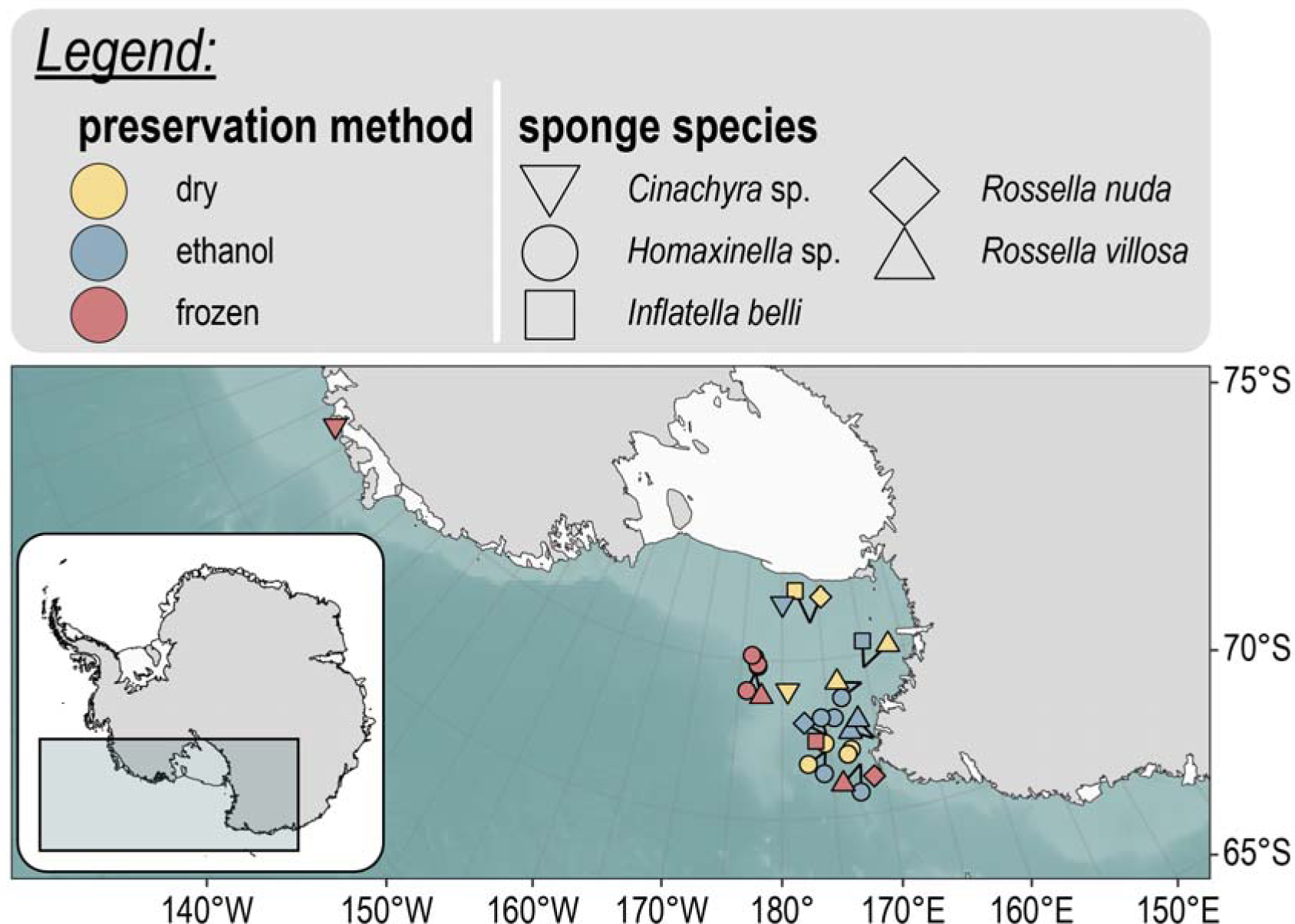
Map of the Ross Sea, Antarctica depicting specimen collection locations. Points are coloured by preservation method: dry (yellow), ethanol (blue), and frozen (red). Point shape is dictated by sponge ID: Cinachyra sp. (inverted triangle), Homaxinella sp. (circle), Inflatella belli (square), Rossella nuda (diamond), and Rossella villosa (triangle).

## 2 MATERIALS & METHODS

### 2.1 Museum specimens

We investigated the potential of extracting heDNA from museum-stored sponge specimens preserved using various techniques, including ethanol submersion, dried, and frozen. Within the NIWA Invertebrate Collection (NIC), dried specimens were initially ethanol preserved followed by dry long-term storage, while frozen specimens are a temporary storage solution until long-term specimen preservation in ethanol. Ten specimens from the Ross Sea (Antarctica) were selected for each preservation technique, covering three orders of Demospongiae (Suberitida; Poecilosclerida; Tetractinellida) and the order Lyssacinosida within the class Hexactinellida (FIG 2; S1 TABLE). Sponges identified as the same genus, and where possible the same species, were processed for each preservation method to limit the potential effect of eDNA accumulation efficiency differences among sponge species (Cai et al. 2022; Brodnicke et al. 2023). To mitigate the potential variation in successful heDNA recovery due to specimen age, we aimed to process specimens collected around a similar collection date. Hence, all specimens included in this experiment were collected and deposited in the NIC between 2004 and 2010, except for the dried *Cinachyra barbata* Sollas, 1886 specimen from 1960, a species for which no dried specimen from the early 2000’s was available.

### 2.2 Laboratory processing of sponge specimens

Five tissue biopsies were collected from each specimen at NIC. Biopsies were transported to the University of Otago’s PCR-free eDNA facilities at Portobello Marine Laboratory (PML) to minimise contamination risk during sample processing. Bench spaces and equipment were sterilised using a 10-minute exposure to 10% bleach dilution (0.5% hypochlorite final concentration) and wiped with ultrapure water (UltraPure^TM^ DNase/RNase-Free Distilled Water, Invitrogen^TM^) before laboratory work (Prince and Andrus 1992). Additionally, negative controls were processed alongside samples during DNA extraction (50 µl ultrapure water) and added as no template controls during qPCR amplification (2 µl ultrapure water). DNA extractions were performed using the Qiagen DNeasy Blood & Tissue Kit (Cat # 69506; Qiagen GmbH, Germany) following the manufacturer’s recommendations, with slight modifications (S2 PROTOCOL). DNA extracts were stored at −20°C until further processing.

Input DNA for qPCR amplification was optimised for each sample using a 10-fold dilution series to identify inhibitors and low-template samples prior to library preparation (Murray et al. 2015). Amplification was carried out in 25 µl duplicate reactions. The qPCR mastermix consisted of 1x SensiMIX SYBR Lo-ROX Mix (Cat # QT625-05; Meridian Bioscience, UK), 0.4 µmol/l of the forward [Fish16SF: 5’-GACCCTATGGAGCTTTAGAC-3’ (Berry et al. 2017)] and reverse [Fish16S2R: 5’-CGCTGTTATCCCTADRGTAACT-3’ (Deagle et al. 2007)] primer (Integrated DNA Technologies, Australia), 2 µl of template DNA, and ultrapure water as required. The thermal profile included an initial denaturation step of 95°C for 10 minutes; followed by 50 cycles of 30 seconds at 95°C, 30 seconds at 54°C, and 45 seconds at 72°C, and a final melt-curve analysis.

Library preparation followed a one-step amplification protocol using fusion primers (Berry et al. 2017). Fusion primers consisted of an Illumina adapter, a modified Illumina sequencing primer, a 6 – 8 bp barcode tag, and the template-specific primer (Fish16SF/Fish16S2R) amplifying a ∼200 bp fragment of the 16S ribosomal RNA gene region. Each sample was amplified in duplicate and assigned a unique barcode combination, whereby forward and reverse barcodes differed from each other in a single sample. The qPCR conditions followed the protocol as described above. Sample duplicates were pooled to reduce stochastic effects from PCR amplification (Leray and Knowlton 2015; Alberdi et al. 2018). Samples were pooled into mini-pools based on end-point qPCR fluorescence, Ct-values, and melt-curve analysis. Mini-pools were visualised using gel electrophoresis to confirm the presence of a single band, and the concentration of mini-pools was measured on Qubit (Cat # Q32854; Qubit^TM^ dsDNA HS Assay Kit, ThermoFisher Scientificc, US). Equimolar pooling produced a single DNA library. Due to differences in cycle number between samples and negative controls, the latter were spiked into the library to allow for optimal library concentration according to Illumina MiSeq® specifications. Size selection was performed using Pippin Prep (Cat # PIP0001; Sage Science, US). The size-selected library was purified using Qiagen’s QIAquick PCR Purification Kit (Cat # 28104; Qiagen GmbH) and quantified using Qubit. Sequencing was performed at the Otago Genomics Facility, University of Otago (New Zealand) on an Illumina MiSeq® instrument using MiSeq reagent kit v2 1×300 bp, with 5 −10% PhiX spiked into the library to minimise issues associated with low-complexity libraries.

### 2.3 Bioinformatic analysis and taxonomy assignment

Prior to bioinformatic processing, raw sequencing files were checked for quality using FastQC *v* 0.11.5 (Andrews 2010). Reverse Illumina adapter sequences, present due to the amplicon size being smaller than the sequencing kit cycle number, were removed from reads using cutadapt *v* 4.1 (Martin 2011) without allowing indels. Reads were demultiplexed and assigned to samples using cutadapt, allowing for two mismatches in the barcode and primer region. The assigned amplicons were filtered using the ‘*--fastq_filter*’ function in VSEARCH *v* 2.13.3 (Rognes et al. 2016) based on a maximum expected error of 1.0, a minimum length of 190 bp, a maximum length of 220 bp, and without allowing the occurrence of ambiguous base calls. The remaining reads were checked for successful quality filtering using FastQC before dereplication (function: ‘*vsearch --derep_fulllength*’). Chimeric sequences were removed and Zero-radius Operational Taxonomic Units (ZOTUs) were generated using the ‘*-unoise3*’ function (Edgar 2016) in USEARCH *v* 11.0.667 (Edgar, 2010). Finally, a frequency table was generated using the ‘*-otutab*’ function in USEARCH.

A custom-curated reference database was generated using CRABS *v* 0.1.5 (Jeunen et al. 2022). The custom curated reference database consisted of sequences downloaded from multiple online repositories using the ‘*db_download*’ function and *in-house* generated barcodes of Southern Ocean fish species (Jeunen et al. 2024) using the ‘*db_import*’ function. Amplicon regions were extracted from sequences through *in silico* PCR analysis (‘*insilico_pcr*’ function) and pairwise global alignments (‘*pga*’ function). Finally, the curated reference database was filtered (function: ‘*seq_cleanup*’) and dereplicated (function: ‘*dereplicate*’). The final reference database was formatted according to IDTAXA specifications (Murali et al. 2018) and used as the reference database (S3 FILE) to train the IDTAXA classifier through five iterations using the ‘*LearnTaxa*’ function in the DECIPHER R package (Wright 2016). Finally, all ZOTU sequences were classified using the ‘*IdTaxa*’ function in DECIPHER, with a threshold of 60% as the cut off value to determine the taxonomic ID level.

After taxonomy assignment, the frequency table underwent final processing before statistical analysis, whereby (i) detections were only kept when reaching a read count higher than the most abundant detection in the summed negative controls, (ii) sequences with a positive detection in the negative controls were deemed true detections in samples when achieving a 10x read count compared to the negative controls, (iii) sequences were removed from the final data set if no taxonomic ID could be obtained for at least the order level, (iv) non-Antarctic taxonomic IDs were removed from the frequency table, (v) artefact sequences were merged with their parent based on taxon-dependent co-occurrence patterns of similar sequences, and (vi) samples not reaching a total abundance of 10,000 reads were removed from the analysis.

### 2.4 Statistical analysis and visualisation

Statistical analyses and visualisations were conducted in R *v* 4.0.5 (R; http://www.R-project.org) unless specified otherwise. Rarefaction curves were generated from the unfiltered frequency table to assess sequencing coverage using the *vegan v* 2.5-7 package (Dixon 2003). Species accumulation curves were drawn for Hill numbers of order q: species richness (q = 0), the exponential of Shannon entropy (q = 1), and the inverse of Simpson concentration (q = 2) to assess replication coverage per specimen using the iNEXT.3D *v* 1.0.1 R package (Chao et al. 2021). Summary statistics on the read count and most abundant taxa were obtained through the phyloseq *v* 1.44.0 (McMurdie and Holmes 2013) and microbiome *v* 1.23.1 R packages. To assess alpha diversity differences among preservation methods, the frequency table was transformed to an incidence-frequency data set. Hill numbers of orders q = 0, 1, and 2 were compared among preservation methods through a one-way ANOVA. Taxonomic diversity estimates for Hill order q = 0 were calculated through inter- and extrapolation in iNEXT.3D (function: ‘*estimate3D*’) to assess the required replication at 90% coverage for each specimen. Significant differences among preservation methods for the required replication were tested through a one-way ANOVA, followed by post hoc Fisher’s LSD (Least Significant Difference). Non-metric MultiDimensional Scaling (NMDS) ordination plots were drawn using the phyloseq function ‘*ordinate*’ to examine beta diversity patterns. Statistical significant differences in beta diversity among preservation methods, sampling methods, sponge IDs, depth, latitude, and longitude were tested through PERMANOVA (function ‘*adonis2*’) and PERMDISP analyses (‘*betadisper*’). Bioinformatic and R scripts and metadata files can be found on the GitHub repository https://github.com/gjeunen/marsden_obj1_preservationMethod. The raw sequence data are deposited onto the NCBI short read archive (SRA) under project number PRJNA1019816.

## 3 RESULTS

### 3.1 High-throughput sequencing results

Demultiplexing of raw sequencing data assigned 10,989,938 sequences to heDNA extracts. Quality filtering and denoising returned a total of 10,680,592 (97.19%) sequences assigned to 153 ZOTUs. Post-processing identified four reads in negative controls, including three reads assigned to ZOTU 2 (*Macrourus* sp.) and one read assigned to ZOTU 8 (*Pleuragramma antarcticum* Boulenger, 1902). Hence, all detections with three reads or lower were discarded from the frequency table, as well as detections with 30 reads or lower and 10 reads or lower for ZOTU 2 and ZOTU 8, respectively. IDTAXA failed to provide a taxonomic ID at the order level for 38 ZOTUs (10,244 reads), which were removed from the analysis. Additionally, three ZOTUs were assigned to temperate taxa and removed from the analysis, including ZOTU 56 (taxonomic ID: Cheilodactylidae; read abundance: 2,203; detections: PMD7b), ZOTU 71 (taxonomic ID: *Helicolenus* sp.; read abundance: 582; detections: PMD7e), and ZOTU 88 (taxonomic ID: *Thyrsites atun* (Euphrasen, 1791); read abundance: 131; detections: PMF3e, PMF9a). After merging artefact sequences, 64 ZOTUs were retained for the final analysis. Nine samples did not obtain a read count of 10,000 sequences, including PMD1d and multiple samples belonging to the sponge genus *Cinachyra spp.* irrespective of the preservation method used (PMD4; PME3; PMF5). Therefore, all samples belonging to genus *Cinachyra* were removed from the analysis. Post-processing of the frequency table retained a total of 9,829,826 (92.03%) reads for statistical analysis (S4 TABLE). Overall, samples achieved sufficient sequencing coverage based on the plateauing of rarefaction curves (S5 FIG) and mean number of reads per sample ± *SD*: 73,357 ± 25,183.

### 3.2 Alpha diversity measurements

Post-processing returned 64 ZOTUs for which a taxonomic ID could be achieved, covering 25 families, fifteen orders, and two classes (FIG 3; S4 TABLE). The Antarctic toothfish (*Dissostichus mawsoni*, Norman, 1937) was the most abundant signal across all samples (sequence ID: ZOTU 1; read count: 2,318,043; proportional abundance: 23.58%), followed by the genus *Macrourus* (sequence ID: ZOTU 2; read count: 1,877,328; proportional abundance: 19.10%) and the Antarctic silverfish (*Pleuragramma antarcticum*; sequence ID: ZOTU 3; read count: 1,856,736; proportional abundance: 18.89%). The Antarctic toothfish was also the most frequently detected species across all samples (detections: 128/134), followed by the Antarctic silverfish (detections: 122/134) and cod icefish in the genus *Trematomus* (sequence ID: ZOTU 4; detections: 106/134).

**FIG 3:**
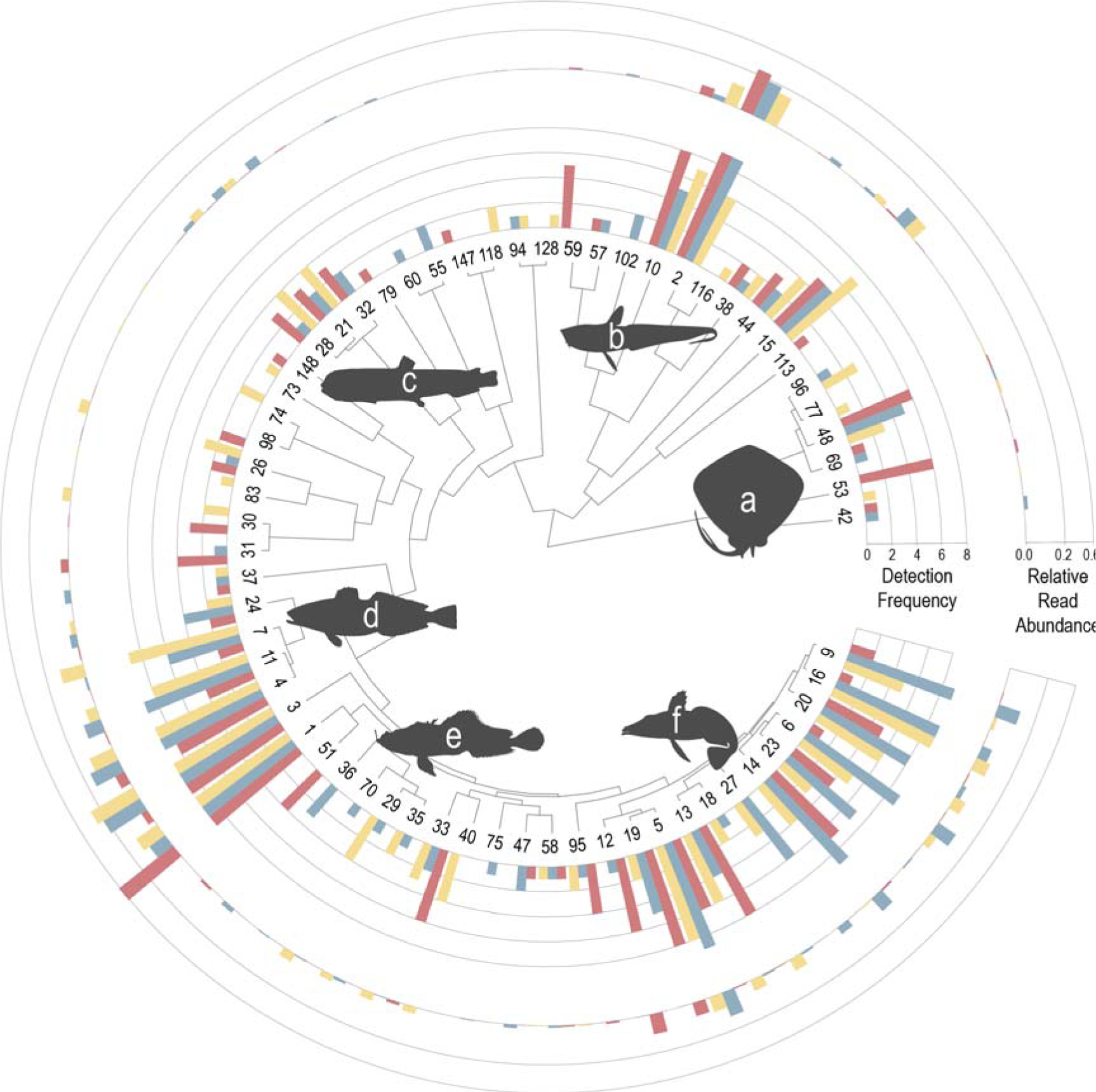
Bayesian phylogenetic tree generated for all 64 ZOTU sequences. Tip labels represent ZOTU number. Taxonomic ID for each ZOTU can be retrieved from SUPPLEMENT 4. Inner bar graph showing the number of detections of each ZOTU sequence within the nine specimens stored dry (yellow), in ethanol (blue), and frozen (red). Outer bar graph showing the relative read abundance of each ZOTU sequence within the nine specimens stored dry (yellow), in ethanol (blue), and frozen (red). Axis for relative read abundance bar graph is reported as square root transformed to increase readability of low-abundant signals. Most frequent and abundant taxonomic groups are represented by silhouettes, including (a) Chondrichthyes, (b) Gadiformes, (c) Bathylagidae, (d) Nototheniidae, (e) Bathydraconidae, and (f) Channichthyidae.

Five DNA extracts per sponge specimen were deemed sufficient to recover most of the fish diversity held within the sponge according to the plateauing of species accumulation curves (S6 FIG). The estimated replication needed to recover 90% of the fish diversity, based on inter- and extrapolation calculations, differed significantly among preservation methods according to a one-way ANOVA (*F*_2,23_ = 3.463, *p* = 0.048*) when removing the outlier sample PMD1 (data from only 4/5 replicates, with 1/5 replicates dropped out). Fisher’s LSD identified frozen specimens (8.840 ± 4.102) to be significantly different from ethanol-stored (5.116 ± 3.224) and dried (5.215 ± 2.579) specimens (FIG 4a). Without removing the outlier sample, no significant differences among preservation methods were observed according to a one-way ANOVA (*F*_2,24_ = 1.893, *p* = 0.173; S7 FIG). Alpha diversity investigations among preservation methods yielded no significant differences across three orders of Hill numbers according to one-way ANOVA (q = 0: *F*_2,24_ = 0.146, *p* = 0.865; q = 1: *F*_2,24_ = 0.237, *p* = 0.791; q = 2: *F*_2,24_ = 0.444, *p* = 0.647; FIG 4b).

**FIG 4:**
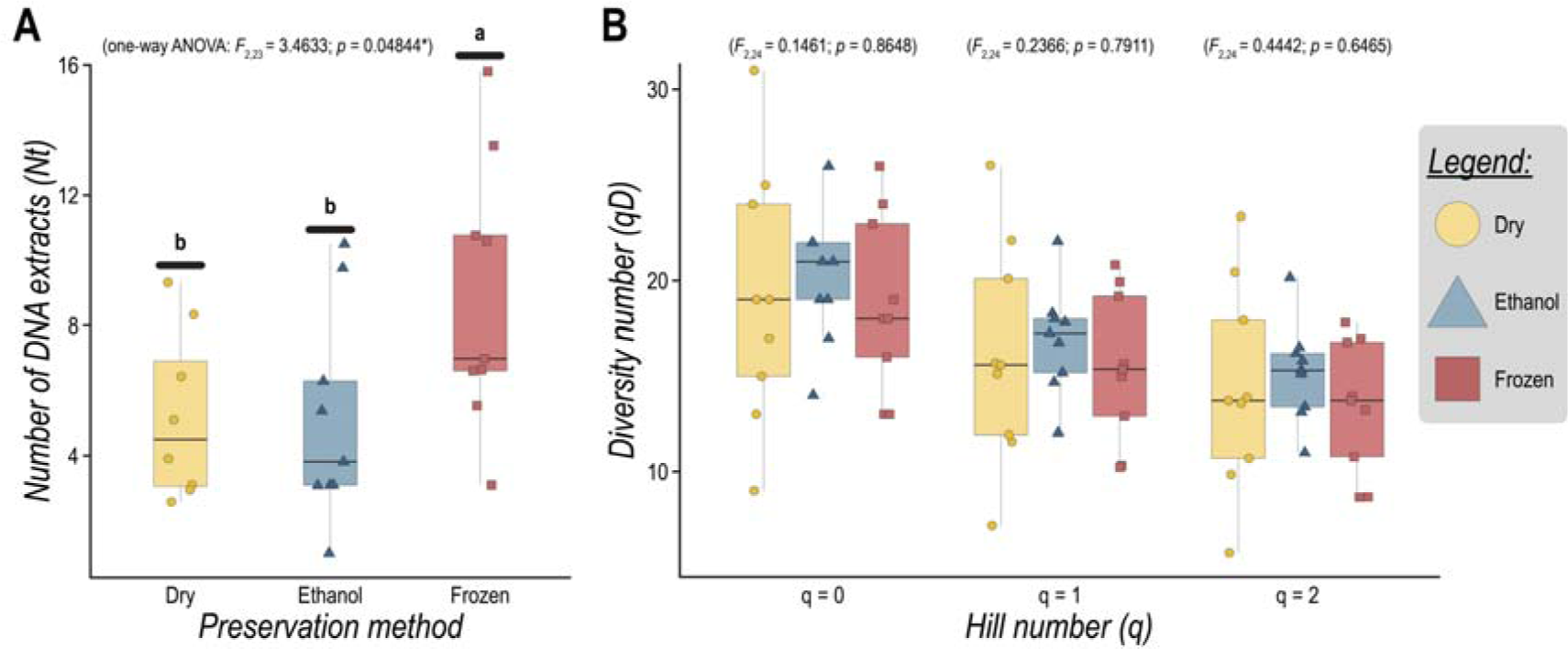
(a) Boxplots depicting the estimated tissue biopsies needed to recover 90% of the fish diversity among the three preservation methods, including dry (yellow), ethanol (blue), and frozen (red). The median is indicated by a black line within each boxplot. Samples are indicated by coloured dots, including circle (dry), triangle (ethanol), and square (frozen). The outlier specimen PMD1 (four out of DNA extracts yielded fish eDNA signals) was removed from the analysis. One-way ANOVA results are presented above the figure. Significant differences among preservation methods, as reported by Fisher’s LSD, are indicated by lower-case letters. (b) Boxplots depicting alpha diversity measurements among the three preservation methods for three orders of Hill numbers, including q = 0 (species richness), q = 1 (exponential of Shannon entropy), and q = 2 (inverse of Simpson concentration). Non-significant one-way ANOVA results are presented above the figure. Colour and shape follows FIG 4a.

### 3.3 Community composition analyses

Significant differences were observed in community composition among preservation methods according to PERMANOVA (*F*_2,26_ = 2.294; R^2^ = 0.129; *p* < 0.005*), while sampling method (*F*_2,26_ = 1.335; R^2^ = 0.075; *p* > 0.1) and sponge ID (*F*_3,26_ = 1.165; R^2^ = 0.098; *p* > 0.1) were found non-significant explanatory variables. However, the largest fraction of the variability in the data set was left unexplained (Residual R^2^ = 0.533). No significant differences in dispersion were detected among preservation methods according to PERMDISP (*F*_2,26_ = 1.936; *p* > 0.1), indicating PERMANOVA significance resulted from different centroid position in ordination space. Historical fish eDNA signal differences were confirmed by ordination analysis (NMDS; Bray-Curtis index; frequency-occurrence transformation; stress = 0.171; FIG 5), whereby the differently coloured preservation methods and filled vs outline for sampling methods highlight the confounding factors of frozen specimens collected through commercial longlining and dried and ethanol-stored specimens collected by scientific trawling.

**FIG 5:**
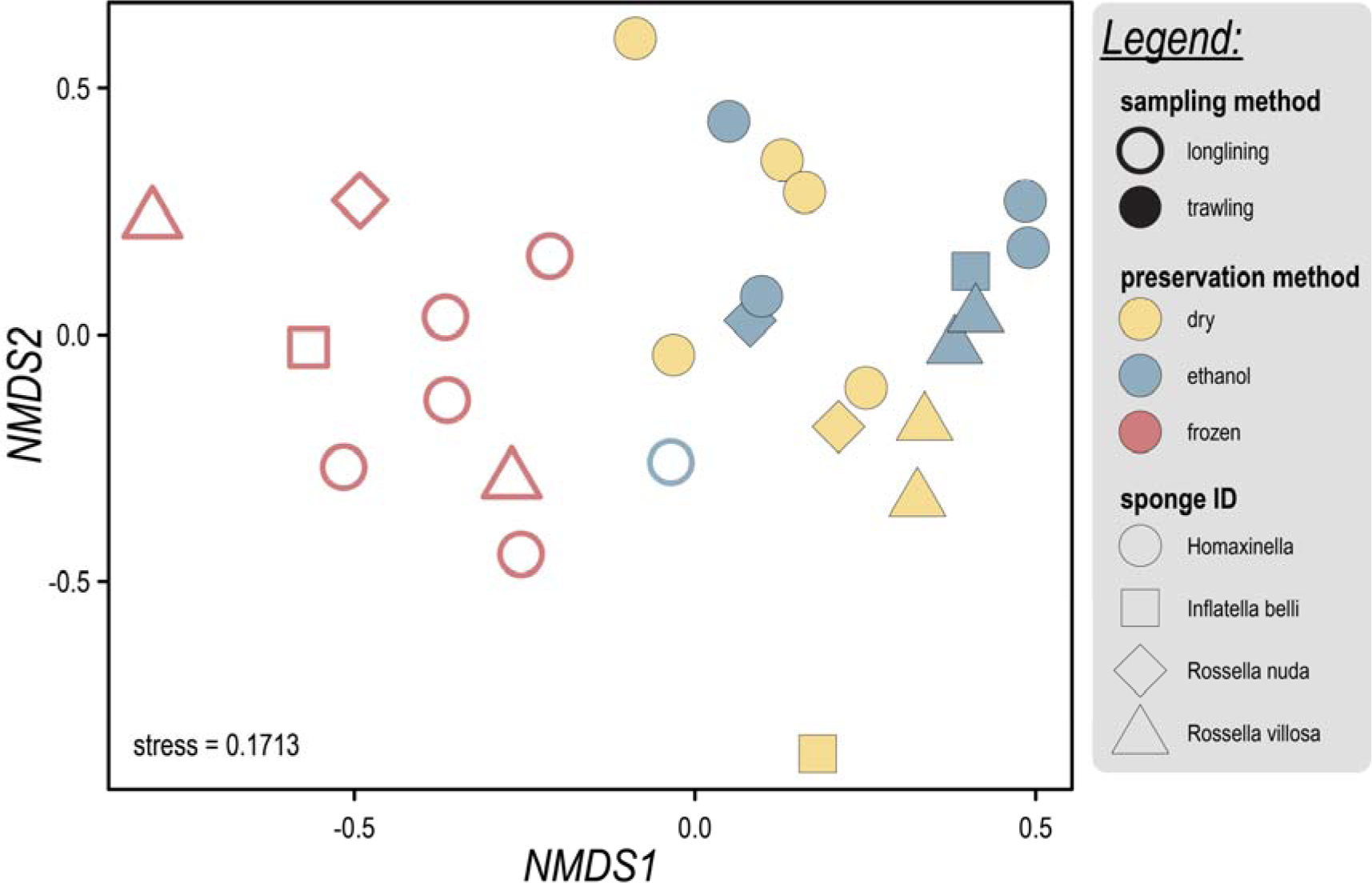
Non-metric multi-dimensional scaling (NMDS) plot depicting similarity in fish community composition based on occurrence frequency (Bray-Curtis index; frequency count). The stress value is reported in the lower left-hand corner. Points are coloured according to preservation method: dry (yellow), ethanol (blue), and frozen (red). Shape is dictated by sponge ID, with Homaxinella sp. represented as circles, Inflatella belli as squares, Rossella nuda as diamonds, and Rossella villosa as triangles. Filled shapes indicate sponge specimens collected through trawling. Outlined shapes indicate sponge specimens collected through longlining.

## 4 DISCUSSION

Environmental DNA biomonitoring has helped increase our understanding of biodiversity and, ultimately, ecosystem functioning (Aglieri et al. 2021; Seymour et al. 2021). Thus far, eDNA has been applied to a range of habitats and locations (Ruppert et al. 2019). Furthermore, the ease of sample collection to monitor biodiversity across the tree of life has made eDNA especially beneficial for remote and logistically-demanding environments that are spatially and temporally under-sampled, such as the Antarctic (Howell et al. 2021; Clarke et al. 2023). Obtaining quantitative spatial and temporal information on Antarctic species is more than ever critical, with the region forecast to see major physical and biological changes in response to climate change and anthropogenic pressures (Chown and Brooks 2019; Convey and Peck 2019). While eDNA has been successfully implemented to monitor contemporary biodiversity patterns of the Antarctic marine biome (Cowart et al. 2018; Clarke et al. 2021; Jeunen et al. 2023b; Liao et al. 2023; Suter et al. 2023), a lack of long-term, quantitative observations limits our understanding of the natural variability in Antarctic ecosystems and complicates future policy making (Howell et al. 2021; Suter et al. 2023). Hence, investigating historical and ancient DNA has the potential to provide the missing information for successful conservation efforts in Antarctica.

In this study, we provide evidence for a widely available but previously untapped resource of historical ecological data that takes advantage of the natural accumulation of eDNA in filter-feeding tissue matrices (Mariani et al. 2019; Neave et al. 2023). Using a targeted metabarcoding approach, we successfully recovered the historical fish eDNA accumulated within Antarctic sponge specimens. Successful DNA extraction from specimens stored using three common preservation techniques, i.e., ethanol submersion, drying, and freezing, increases the number of specimens available for analysis. With vast numbers of marine sponges having been gathered globally since the earliest scientific voyages (Wulff 2016), these archived specimens provide unique ecosystem time capsules through which we can reconstruct historical biodiversity patterns and provide essential knowledge for current conservation efforts (Revéret et al. 2023).

We were able to identify a diverse profile of Actinopterygii and Chondrichthyes from Antarctic sponge specimens, irrespective of the preservation method used. The Antarctic fish community constituted 64 taxa ranging from Nototheniidae (cod icefishes) and Channichthyidae (icefishes), to Bathydraconidae (Antarctic dragonfishes), all of which are known to occur in the Ross Sea according to Antarctic toothfish bycatch records (Pinkerton and Bradford-Grieve 2014; Jeunen et al. 2024). Interestingly, one notably absent taxonomic group, besides two signals of *Gymnoscopelus* sp., from sponge specimens collected in deeper waters are the myctophids, the most diverse and abundant group of mesopelagic fishes globally, including in the Southern Ocean (Duhamel et al. 2014; Woods et al. 2023; Vasiliadis et al. 2024). The lack of myctophid detection could potentially have stemmed from their occupancy of the mesopelagic zone (Catul et al. 2011; Christiansen et al. 2018).

The vertical distance between myctophids and benthic sponges is known to influence eDNA metabarcoding detection results (Jeunen et al. 2019b). Additionally, multiple mismatches at the 3’ end of the forward PCR primer-binding region (S8 FIG) could have significantly reduced the amplification efficiency for this taxonomic group, resulting in false-negative detections (Stadhouders et al. 2010). While universal metabarcoding approaches have been reported to be an inefficient solution due to the co-amplification of sponge host DNA (Jeunen et al. 2023b), a multi-marker targeted metabarcoding approach has previously been proposed for aquatic eDNA research to increase species detection accuracy and reduce the impact of amplification bias (McElroy et al. 2020).

Our results provide evidence for the importance of accurate metadata to interpret observed biodiversity patterns and gauge the potential impact of biases in species detection from eDNA metabarcoding. For example, the taxonomic group to which a sponge belongs has been identified in previous research (Cai et al. 2022; Brodnicke et al. 2023), as well as here, to impact eDNA detection success. In our study, specimens from the genus *Cinachyra* failed to reliably amplify fish eDNA signals, irrespective of the preservation technique used to store the specimens. Additionally, a significant difference in the reported fish community was observed among preservation methods. This difference, however, could have originated from the confounding factors of collection location and method. Namely, frozen specimens were collected by commercial Antarctic toothfish longlining vessels located further offshore compared to dried and ethanol-stored specimens collected by scientific trawling along the Ross Sea coastline.

Thus far, contemporary sponge eDNA research has focused on single-tissue biopsies for eDNA signal detection (Mariani et al. 2019; Cai et al. 2022; Brodnicke et al. 2023; Neave et al. 2023). However, replicate biopsies collected from a single sponge specimen combined with rarefaction and extrapolation of species diversity identified the need to collect between five (dried and ethanol-stored) and nine (frozen) biopsies per specimen to confidently detect 90% of the fish diversity held within marine sponges. While multiple tissue biopsies from each sponge increases the overall cost of the project and may not be possible for small and/or rare specimens in collections, replication enables data transformation to frequency-occurrence (Chao et al. 2021), thereby providing semi-quantitative, i.e., incidence-based, data and expand upon the statistical analyses able to be conducted (Alberdi and Gilbert 2019). The need for increased replication likely stems from a lack of understanding about the process of eDNA accumulation in the sponge tissue matrix. Further research into laboratory protocol development to efficiently extract eDNA from sponge tissues (Harper et al. 2023), as well as gaining a better understanding of eDNA accumulation by sponges (Cai et al. 2022), are essential to progress the applicability of sponges as natural eDNA samplers. Our results show significant differences in estimated replication between treatments, with frozen samples requiring increased tissue biopsies to reliably detect 90% of the fish diversity within a specimen compared to dried and ethanol-submerged specimens. The significant difference in the required replication could have been induced by the highly dominant signal of *D. mawsoni*, the target fish of the longlining fishing vessels from which frozen specimens were collected, thereby reducing the detection probability of the remaining low-abundant fish eDNA signals (Ficetola et al. 2015; Bylemans et al. 2019; Rojahn et al. 2021).

The challenge in verifying species detection became evident from the presence of temperate fish species in our dataset. All temperate fish species were conspicuously absent in the negative control samples, thus unlikely to be a result from internal lab contamination. The power and sensitivity of present-day molecular approaches require high standards to minimise the risk of DNA contamination in the field and throughout curation and laboratory handling (Goldberg et al. 2016; Llamas et al. 2017) Processing ancient and historical specimens, most of which were not collected nor handled for molecular analysis purposes throughout the time stored in scientific collections, increases the risk of DNA contaminants being incorporated into the specimens through, for example, (i) cross-contamination from handling multiple specimens without bench-space and equipment sterilisation, or (ii) transferring specimens and fixatives between collection lots (Knapp et al. 2012; Cowart et al. 2022). For ancient DNA shotgun sequencing approaches, DNA damage profiles can be assessed to identify modern DNA contaminants (Seersholm et al. 2016). However, when utilising historical metabarcoding techniques, DNA damage profiles cannot be successfully implemented for contaminant identification (Piper et al. 2019). Within eDNA metabarcoding and microbiome research, removal of contaminants has been largely based on abundance filtering (Li et al. 2018), detection frequency filtering (Evans et al. 2017), and removal of non-target species (Alberdi et al. 2018), as employed in this study.

The selection of preservation methods included in this study was determined by identifying the techniques with the highest number of sponge specimens within NIC. Exploring additional common curation methods, such as formalin fixation (Srinivasan et al. 2002; Hykin et al. 2015), will further increase the pool of available specimens for historical eDNA research. Genetic and genomic investigations utilising formalin-fixed museum specimens have been challenging in the past, since formaldehyde reduces DNA integrity and produces sequence artefacts by inducing numerous molecular lesions, such as strand breaks, base misincorporation, and intra- and intermolecular cross-linking (Williams et al. 1999; Srinivasan et al. 2002; Do and Dobrovic 2015). However, recent advances in whole-genome sequencing of formalin-fixed paraffin-embedded (FFPE) archival tissues (Stiller et al. 2016; Robbe et al. 2018) and formalin-fixed museum specimens (Hahn et al. 2021) provides a tantalising prospect to explore formalin-fixed sponge specimens for historical eDNA research, which we will seek to undertake in future studies.

To fully utilise the power of this novel historical resource, we propose three future research avenues. First, while our results provide evidence for successful heDNA extraction following multiple preservation techniques, investigations into optimal storage methods and associated biases require specimens to be divided and preserved in various ways (Spens et al. 2016). Such information will guide scientists in choosing optimal specimens for heDNA research (Hahn et al. 2021) and set storage standards for building future resources. Second, to minimise the number of covariates in this study, we aimed to incorporate specimens from a similar collection date range in the experiment. Further investigations into the effect of specimen age for each preservation technique would provide useful information on the utility for older specimens. While the dried *C. barbata* specimen from 1960 failed to amplify historical fish eDNA signals, the result was most likely influenced by sponge taxonomy rather than age, as we successfully amplified and analysed the oldest Antarctic sponge specimen stored in ethanol (collection date: 1958) at the NIC for fish eDNA signals (Jeunen, *pers. comm.*). Third, museum specimens are precious but finite resources for scientific research (Hahn et al. 2020). Therefore, minimising the destruction of valuable voucher specimens is essential and will require the use of optimised wet lab protocols, as well as investigations into non-destructive DNA extraction approaches, such as direct heDNA extraction from preservative medium rather than tissue biopsies (Rohland et al. 2004; Shokralla et al. 2010).

## 5 CONCLUSION

Marine environments and species have been exploited throughout human history, leading to entire ecosystem modification, habitat degradation, and multiple species extinctions. Therefore, mitigation and restoration of degraded marine systems is of top global economic, ecological, and cultural importance. However, successful remediation requires detailed knowledge of how these ecosystems have altered over time. Currently, the extent and speed of ecological change in the marine domain have rarely been quantified because long-term ecological records are scarce and accurate historical data are difficult and expensive to obtain. In this experiment, we provide evidence for using the historical eDNA trapped within taxonomic collection sponge specimens as a novel ecological record source to investigate historical biodiversity patterns at a previously unattainable temporal and spatial scale. The successful recovery of historical eDNA from sponge specimens stored using various preservation techniques significantly broadens the pool of specimens to be included in this type of research. Future investigations into the impact of additional preservation techniques such as formalin-fixation, as well as specimen age, and collection method are essential to fully utilise this novel methodology.

## ACKNOWLEDGEMENTS

A New Zealand Royal Society Te Apārangi Marsden Fast-Start (MFP-UOO2116), a University of Otago Research Grant, and the Ministry of Business, Innovation, and Employment Antarctic Science Platform (MBIE ANTA1801) funded the cost for this project. We thank the NIWA Invertebrate Collection (NIC) for allowing us to utilise the collection for this work and for their assistance in subsampling the specimens used in this experiment. Antarctic specimens were collected from a variety of research projects including from CCAMLR fishery observers, the TAN0402 ‘BioRoss’ survey funded by the former NZ Ministry of Fisheries (MFish), and TAN0802 IPY-CAML survey, a collaborative project by Land Information New Zealand (LINZ), MFish, Ministry of Foreign Affairs and Trade, Antarctica New Zealand, Te papa, NIWA and New Zealand universities. We thank Michelle Kelly for identifying NIC specimens based on spicule analysis and morphological assessments conducted between 2007 and 2010.

## 6 AUTHOR CONTRIBUTIONS

The study design was conceptualised by GJJ, SM, SM, ML, JLS and NJG. Specimen biopsies were collected by GJJ and SM. Laboratory work was performed by GJJ, JT, and SF. Sequencing was conducted by MZ. The bioinformatic analysis was conducted by GJJ. GJJ performed the statistical analysis, with input from SM, SM, ML, GAD, and NJG. GJJ wrote the manuscript with significant input from ML, SM, MK, BDV and NJG. All co-authors contributed to the writing of the manuscript and approve of the submission.

## 7 DATA AVAILABILITY STATEMENT

Bioinformatic and R scripts, as well as metadata files can be found on the GitHub repository https://github.com/gjeunen/marsden_obj1_preservationMethod. The raw sequence data are deposited onto the NCBI short read archive (SRA) under project number PRJNA1019816.

## 8 SUPPORTING INFORMATION

**S1 Table. NIWA Invertebrate Collection sponge specimen metadata.**

**S2 Protocol. Historical eDNA extraction protocol from museum-stored sponges.**

**S3 File. IDTAXA reference database for taxonomy assignment. S4 Table. ZOTU frequency table.**

**S5 Fig. Rarefaction curves.**

**S6 Fig. Species accumulation curves.**

**S7 Fig. Estimated replication analysis without removal of outlier. S8 Fig. *In silico* PCR analysis of myctophids.**

## REFERENCES

Aglieri G, Baillie C, Mariani S, Cattano C, Calò A, Turco G, et al. Environmental DNA effectively captures functional diversity of coastal fish communities. Mol Ecol. 2021;30:3127–39.

Alberdi A, Aizpurua O, Gilbert MTP, Bohmann K. Scrutinizing key steps for reliable metabarcoding of environmental samples. Methods in Ecology and Evolution [Internet]. 2018;9(1):134–47. Available from: 10.1111/2041-210X.12849

Alberdi A, Gilbert MTP. A guide to the application of Hill numbers to DNA-based diversity analyses. Molecular Ecology Resources [Internet]. 2019 Jul 1;19(4):804–17. Available from: 10.1111/1755-0998.13014

Andrews S. FastQC: a quality control tool for high throughput sequence data. Babraham Bioinformatics, Babraham Institute, Cambridge, United Kingdom; 2010.

Aronson RB, Thatje S, McClintock JB, Hughes KA. Anthropogenic impacts on marine ecosystems in Antarctica. Annals of the New York Academy of Sciences. 2011;1223(1):82–107.

Berry TE, Osterrieder SK, Murray DC, Coghlan ML, Richardson AJ, Grealy AK, et al. DNA metabarcoding for diet analysis and biodiversity: A case study using the endangered Australian sea lion (Neophoca cinerea). Ecology and Evolution [Internet]. 2017;7(14):5435–53. Available from: 10.1002/ece3.3123

Brodnicke OB, Meyer HK, Busch K, Xavier JR, Knudsen SW, Møller PR, et al. Deep-sea sponge derived environmental DNA analysis reveals demersal fish biodiversity of a remote Arctic ecosystem. Environmental DNA [Internet]. 2023 Nov 1;5(6):1405–17. Available from: 10.1002/edn3.451

Bylemans J, Gleeson DM, Duncan RP, Hardy CM, Furlan EM. A performance evaluation of targeted eDNA and eDNA metabarcoding analyses for freshwater fishes. Environmental DNA. 2019;1(4):402–14.

Cai W, Harper LR, Neave EF, Shum P, Craggs J, Arias MB, et al. Environmental DNA persistence and fish detection in captive sponges. Molecular Ecology Resources [Internet]. 2022;n/a(n/a). Available from: 10.1111/1755-0998.13677

Catul V, Gauns M, Karuppasamy PK. A review on mesopelagic fishes belonging to family Myctophidae. Rev Fish Biol Fish [Internet]. 2011 Sep 1;21(3):339–54. Available from: 10.1007/s11160-010-9176-4

Cecchetto M, Cesare AD, Eckert E, Fassio G, Fontaneto D, Moro I, et al. Antarctic coastal nanoplankton dynamics revealed by metabarcoding of desalination plant filters: Detection of short-term events and implications for routine monitoring. Science of the Total Environment [Internet]. 2021 Feb 25;757:143809. Available from: https://www.sciencedirect.com/science/article/pii/S004896972037340X

Chao A, Henderson PA, Chiu CH, Moyes F, Hu KH, Dornelas M, et al. Measuring temporal change in alpha diversity: A framework integrating taxonomic, phylogenetic and functional diversity and the iNEXT.3D standardization. Methods in Ecology and Evolution [Internet]. 2021 Oct 1;12(10):1926–40. Available from: 10.1111/2041-210X.13682

Chown SL, Brooks CM. The State and Future of Antarctic Environments in a Global Context. Annual Review of Environment and Resources [Internet]. 2019 Oct 17;44(1):1–30. Available from: 10.1146/annurev-environ-101718-033236

Christiansen H, Dettai A, Heindler FM, Collins MA, Duhamel G, Hautecoeur M, et al. Diversity of Mesopelagic Fishes in the Southern Ocean - A Phylogeographic Perspective Using DNA Barcoding. Frontiers in Ecology and Evolution [Internet]. 2018;6. Available from: https://www.frontiersin.org/articles/10.3389/fevo.2018.00120;

Clarke LJ, Shaw JD, Suter L, Atalah J, Bergstrom DM, Biersma EM, et al. An expert-driven framework for applying eDNA tools to improve biosecurity in the Antarctic. Management of Biological Invasions. 2023;

Clarke LJ, Suter L, Deagle BE, Polanowski AM, Terauds A, Johnstone GJ, et al. Environmental DNA metabarcoding for monitoring metazoan biodiversity in Antarctic nearshore ecosystems. Pochon“] [”Xavier, editor. PeerJ [Internet]. 2021;9:e12458. Available from: 10.7717/peerj.12458

Convey P, Peck LS. Antarctic environmental change and biological responses. Science Advances [Internet]. 2019;5(11):eaaz0888. Available from: 10.1126/sciadv.aaz0888

Cowart DA, Murphy KR, Cheng CHC. Environmental DNA from Marine Waters and Substrates: Protocols for Sampling and eDNA Extraction. In New York, NY: Springer US; 2022. p. 225–51. (Methods in Molecular Biology; vol. 2498). Available from: 10.1007/978-1-0716-2313-8_11

Cowart DA, Murphy KR, Cheng CHCHC. Metagenomic sequencing of environmental DNA reveals marine faunal assemblages from the West Antarctic Peninsula. Marine Genomics [Internet]. 2018;37(September 2017):148–60. Available from: https://www.sciencedirect.com/science/article/pii/S1874778717302581 10.1016/j.margen.2017.11.003

Deagle BE, Gales NJ, Evans K, Jarman SN, Robinson S, Trebilco R, et al. Studying Seabird Diet through Genetic Analysis of Faeces: A Case Study on Macaroni Penguins (Eudyptes chrysolophus). PLOS ONE [Internet]. 2007;2(9):e831. Available from: 10.1371/journal.pone.0000831

Dixon P. VEGAN, a package of R functions for community ecology. Journal of Vegetation Science. 2003;14(6):927–30.

Do H, Dobrovic A. Sequence Artifacts in DNA from Formalin-Fixed Tissues: Causes and Strategies for Minimization. Clin Chem [Internet]. 2015;61(1):64–71. Available from: 10.1373/clinchem.2014.223040

Duhamel G, Hulley PA, Causse R, Koubbi P, Vacchi M, Pruvost P, et al. Biogeographic patterns of fish. In 2014.

Edgar RC. UNOISE2: improved error-correction for Illumina 16S and ITS amplicon sequencing. bioRxiv [Internet]. 2016;81257. Available from: http://biorxiv.org/content/early/2016/10/15/081257.abstract

Evans NT, Li Y, Renshaw MA, Olds BP, Deiner K, Turner CR, et al. Fish community assessment with eDNA metabarcoding: effects of sampling design and bioinformatic filtering. Canadian Journal of Fisheries and Aquatic Sciences [Internet]. 2017 Sep 1;74(9):1362–74. Available from: 10.1139/cjfas-2016-0306

Ficetola GF, Pansu J, Bonin A, Coissac E, Giguet-Covex C, Barba MD, et al. Replication levels, false presences and the estimation of the presence/absence from eDNA metabarcoding data. Molecular Ecology Resources [Internet]. 2015 May 1;15(3):543–56. Available from: 10.1111/1755-0998.12338

Finnegan S, Anderson SC, Harnik PG, Simpson C, Tittensor DP, Byrnes JE, et al. Paleontological baselines for evaluating extinction risk in the modern oceans. Science [Internet]. 2015;348(6234):567–70. Available from: http://science.sciencemag.org/content/348/6234/567.abstract

Finney BP, Gregory-Eaves I, Douglas MSV, Smol JP. Fisheries productivity in the northeastern Pacific Ocean over the past 2,200 years. Nature [Internet]. 2002;416(6882):729–33. Available from: 10.1038/416729a

Ghiglione C, Alvaro MC, Cecchetto M, Canese S, Downey R, Guzzi A, et al. Porifera collection of the Italian National Antarctic Museum (MNA), with an updated checklist from Terra Nova Bay (Ross Sea). ZooKeys [Internet]. 2018;758:137–56. Available from: 10.3897/zookeys.758.23485

Godefroy N, Goff EL, Martinand-Mari C, Belkhir K, Vacelet J, Baghdiguian S. Sponge digestive system diversity and evolution: filter feeding to carnivory. Cell and Tissue Research. 2019;377:341–51.

Goldberg CS, Turner CR, Deiner K, Klymus KE, Thomsen PF, Murphy MA, et al. Critical considerations for the application of environmental DNA methods to detect aquatic species. [“Gilbert, M.”], editors. Methods in Ecology and Evolution [Internet]. 2016;7(11):1299–307. Available from: http://doi.wiley.com/10.1111/2041-210X.12595

Hahn EE, Alexander MR, Grealy A, Stiller J, Gardiner DM, Holleley CE. Unlocking inaccessible historical genomes preserved in formalin. Molecular Ecology Resources [Internet]. 2021;n/a(n/a). Available from: 10.1111/1755-0998.13505

Hahn EE, Grealy A, Alexander M, Holleley CE. Museum Epigenomics: Charting the Future by Unlocking the Past. Trends in Ecology & Evolution. 2020;

Harnik PG, Lotze HK, Anderson SC, Finkel ZV, Finnegan S, Lindberg DR, et al. Extinctions in ancient and modern seas. Trends in Ecology & Evolution [Internet]. 2012;27(11):608–17. Available from: http://www.sciencedirect.com/science/article/pii/S0169534712001711

Harper LR, Neave EF, Sellers GS, Cunnington AV, Arias MB, Craggs J, et al. Optimized DNA isolation from marine sponges for natural sampler DNA metabarcoding. Environmental DNA [Internet]. 2023;n/a(n/a). Available from: 10.1002/edn3.392

Hoegh-Guldberg O, Bruno JF. The Impact of Climate Change on the World’s Marine Ecosystems. Science [Internet]. 2010;328(5985):1523–8. Available from: http://science.sciencemag.org/content/328/5985/1523.abstract

Howell L, LaRue M, Flanagan SP. Environmental DNA as a tool for monitoring Antarctic vertebrates. New Zealand Journal of Zoology [Internet]. 2021 Oct 2;48(3–4):245–62. Available from: 10.1080/03014223.2021.1900299

Hykin SM, Bi K, McGuire JA. Fixing Formalin: A Method to Recover Genomic-Scale DNA Sequence Data from Formalin-Fixed Museum Specimens Using High-Throughput Sequencing. PLOS ONE [Internet]. 2015;10(10):e0141579. Available from: 10.1371/journal.pone.0141579

Iyavoo S, Hadi S, Goodwin W. Evaluation of five preservation methods for recovery of DNA from bone. The 28th Congress of the International Society for Forensic Genetics [Internet]. 2019 Dec 1;7(1):200–2. Available from: https://www.sciencedirect.com/science/article/pii/S1875176819301337

Jensen MR, Sigsgaard EE, Ávila M de P, Agersnap S, Brenner-Larsen W, Sengupta ME, et al. Short-term temporal variation of coastal marine eDNA. Environmental DNA [Internet]. 2022;n/a(n/a). Available from: 10.1002/edn3.285

Jeunen GJ, Cane JS, Ferreira S, Strano F, Ammon U von, Cross H, et al. Assessing the utility of marine filter feeders for environmental DNA (eDNA) biodiversity monitoring. Molecular Ecology Resources [Internet]. 2023a May 1;23(4):771–86. Available from: 10.1111/1755-0998.13754

Jeunen GJ, Dowle E, Edgecombe J, Ammon U von, Gemmell N, Cross H. CRABS--A software program to generate curated reference databases for metabarcoding sequencing data. Molecular Ecology Resources. 2022;

Jeunen GJ, Knapp M, Spencer HG, Lamare MD, Taylor HR, Stat M, et al. Environmental DNA (eDNA) metabarcoding reveals strong discrimination among diverse marine habitats connected by water movement. Molecular Ecology Resources. 2019a;19(2):426–38.

Jeunen GJ, Lamare M, Cummings V, Treece J, Ferreira S, Massuger J, et al. Unveiling the hidden diversity of marine eukaryotes in the Ross Sea: A comparative analysis of seawater and sponge eDNA surveys. Environmental DNA [Internet]. 2023b Dec 6;n/a(n/a). Available from: 10.1002/edn3.500

Jeunen GJ, Lamare M, Devine J, Mariani S, Mills S, Treece J, et al. Characterizing Antarctic fish assemblages using eDNA obtained from marine sponge bycatch specimens. Reviews in Fish Biology and Fisheries [Internet]. 2024 Mar 1;34(1):221–38. Available from: 10.1007/s11160-023-09805-3

Jeunen GJ, Lamare MD, Knapp M, Spencer HG, Taylor HR, Stat M, et al. Water stratification in the marine biome restricts vertical environmental DNA (eDNA) signal dispersal. Environmental DNA [Internet]. 2019b;. Available from: https://onlinelibrary.wiley.com/doi/abs/10.1002/edn3.49

Junk I, Schmitt N, Krehenwinkel H. Tracking climate-change-induced biological invasions by metabarcoding archived natural eDNA samplers. Curr Biol [Internet]. 2023 Sep 25;33(18):R943–4. Available from: https://www.sciencedirect.com/science/article/pii/S0960982223009752

Kidwell SM. Biology in the Anthropocene: Challenges and insights from young fossil records. Proceedings of the National Academy of Sciences [Internet]. 2015;112(16):4922–9. Available from: http://www.pnas.org/content/112/16/4922.abstract

Kittinger JN, McClenachan L, Gedan KB, Blight LK. Marine historical ecology in conservation: Applying the past to manage for the future. Univ of California Press; 2015.

Knapp M, Clarke AC, Horsburgh KA, Matisoo-Smith EA. Setting the stage – Building and working in an ancient DNA laboratory. Special Issue: Ancient DNA [Internet]. 2012 Jan 20;194(1):3–6. Available from: https://www.sciencedirect.com/science/article/pii/S0940960211000732

Kuwae M, Tamai H, Doi H, Sakata MK, Minamoto T, Suzuki Y. Sedimentary DNA tracks decadal-centennial changes in fish abundance. Communications biology. 2020;3(1):1–12.

Leray M, Knowlton N. DNA barcoding and metabarcoding of standardized samples reveal patterns of marine benthic diversity. Proceedings of the National Academy of Sciences of the United States of America [Internet]. 2015;112(7):2076–81. Available from: http://www.ncbi.nlm.nih.gov/pmc/articles/PMC4343139/

Li J, Handley LJL, Read DS, Hänfling B. The effect of filtration method on the efficiency of environmental DNA capture and quantification via metabarcoding. Molecular Ecology Resources [Internet]. 2018;18(5):1102–14. Available from: 10.1111/1755-0998.12899

Liao Y, Miao X, Wang R, Zhang R, Li H, Lin L. First pelagic fish biodiversity assessment of Cosmonaut Sea based on environmental DNA. Mar Environ Res [Internet]. 2023 Nov 1;192:106225. Available from: https://www.sciencedirect.com/science/article/pii/S0141113623003537

Llamas B, Valverde G, Fehren-Schmitz L, Weyrich LS, Cooper A, Haak W. From the field to the laboratory: Controlling DNA contamination in human ancient DNA research in the high-throughput sequencing era. STAR: Science & Technology of Archaeological Research [Internet]. 2017 Jan 1;3(1):1–14. Available from: 10.1080/20548923.2016.1258824

Lotze HK, Worm B. Historical baselines for large marine animals. Trends in Ecology & Evolution [Internet]. 2009;24(5):254–62. Available from: http://www.sciencedirect.com/science/article/pii/S0169534709000482

Lynggaard C, Bertelsen MF, Jensen CV, Johnson MS, Frøslev TG, Olsen MT, et al. Airborne environmental DNA for terrestrial vertebrate community monitoring. Current Biology [Internet]. 2022;32(3):701–707.e5. Available from: https://www.sciencedirect.com/science/article/pii/S0960982221016900

Maiello G, Talarico L, Carpentieri P, Angelis FD, Franceschini S, Harper LR, et al. Little samplers, big fleet: eDNA metabarcoding from commercial trawlers enhances ocean monitoring. Fisheries Research [Internet]. 2022;249:106259. Available from: https://www.sciencedirect.com/science/article/pii/S0165783622000364

Mariani S, Baillie C, Colosimo G, Riesgo A. Sponges as natural environmental DNA samplers. Current Biology [Internet]. 2019;29(11):R401–2. Available from: http://www.sciencedirect.com/science/article/pii/S0960982219304294

Martin M. Cutadapt removes adapter sequences from high-throughput sequencing reads. EMBnet journal. 2011;17(1):10–2.

Martínková N, Searle JB. Amplification success rate of DNA from museum skin collections: a case study of stoats from 18 museums. Molecular Ecology Notes [Internet]. 2006 Dec 1;6(4):1014–7. Available from: 10.1111/j.1471-8286.2006.01482.x

McElroy ME, Dressler TL, Titcomb GC, Wilson EA, Deiner K, Dudley TL, et al. Calibrating Environmental DNA Metabarcoding to Conventional Surveys for Measuring Fish Species Richness. Frontiers in Ecology and Evolution [Internet]. 2020;8. Available from: https://www.frontiersin.org/articles/10.3389/fevo.2020.00276;

McMurdie PJ, Holmes S. phyloseq: An R Package for Reproducible Interactive Analysis and Graphics of Microbiome Census Data. PLOS ONE [Internet]. 2013 Apr 22;8(4):e61217. Available from: 10.1371/journal.pone.0061217

Minamoto T, Fukuda M, Katsuhara KR, Fujiwara A, Hidaka S, Yamamoto S, et al. Environmental DNA reflects spatial and temporal jellyfish distribution. PLOS ONE [Internet]. 2017;12(2):1–15. Available from: 10.1371/journal.pone.0173073

Moreau CS, Wray BD, Czekanski-Moir JE, Rubin BER. DNA preservation: a test of commonly used preservatives for insects. Invertebrate Systematics [Internet]. 2013 Mar 1;27(1):81–6. Available from: 10.1071/IS12067

Murali A, Bhargava A, Wright ES. IDTAXA: a novel approach for accurate taxonomic classification of microbiome sequences. Microbiome. 2018;6(1):1–14.

Murray DC, Coghlan ML, Bunce M. From Benchtop to Desktop: Important Considerations when Designing Amplicon Sequencing Workflows. PLOS ONE [Internet]. 2015;10(4):1–21. Available from: 10.1371/journal.pone.0124671

Nagy ZT. A hands-on overview of tissue preservation methods for molecular genetic analyses. Org Divers Evol [Internet]. 2010 Mar 1;10(1):91–105. Available from: 10.1007/s13127-010-0012-4

Neave EF, Cai W, Arias MB, Harper LR, Riesgo A, Mariani S. Trapped DNA fragments in marine sponge specimens unveil north Atlantic deep-sea fish diversity. Proceedings of the Royal Society B. 2023;2023–03.

Nishiguchi MK, Doukakis P, Egan M, Kizirian D, Phillips A, Prendini L, et al. DNA isolation procedures. Springer; 2002.

Norris RD, Turner SK, Hull PM, Ridgwell A. Marine Ecosystem Responses to Cenozoic Global Change. Science [Internet]. 2013;341(6145):492–8. Available from: http://science.sciencemag.org/content/341/6145/492.abstract

O’Donnell JL, Kelly RP, Shelton AO, Samhouri JF, Lowell NC, Williams GD. Spatial distribution of environmental DNA in a nearshore marine habitat. [“Johnson, Magnus”], editors. PeerJ [Internet]. 2017;5:e3044. Available from: 10.7717/peerj.3044

Olmedo-Rojas P, Jeunen GJ, Lamare M, Turnbull J, Terauds A, Gemmell N, et al. Soil environmental DNA metabarcoding in low-biomass regions requires protocol optimization: a case study in Antarctica. Antarctic Science [Internet]. 2023;35(1):15–30. Available from: https://www.cambridge.org/core/product/EFE7DB6E075AA07A428D81CFCB8F41FB

Parducci L, Nota K, Wood J. Reconstructing Past Vegetation Communities Using Ancient DNA from Lake Sediments BT - Paleogenomics: Genome-Scale Analysis of Ancient DNA. In Cham: Springer International Publishing; 2019. p. 163–87.

Parkinson CL. A 40-y record reveals gradual Antarctic sea ice increases followed by decreases at rates far exceeding the rates seen in the Arctic. Proceedings of the National Academy of Sciences [Internet]. 2019;116(29):14414 LP – 14423. Available from: http://www.pnas.org/content/116/29/14414.abstract

Pauly D, Zeller D. Catch reconstructions reveal that global marine fisheries catches are higher than reported and declining. Nature Communications [Internet]. 2016;7(1):10244. Available from: 10.1038/ncomms10244

Pinkerton MH, Bradford-Grieve JM. Characterizing foodweb structure to identify potential ecosystem effects of fishing in the Ross Sea, Antarctica. ICES Journal of Marine Science. 2014;71(7):1542–53.

Piper AM, Batovska J, Cogan NOI, Weiss J, Cunningham JP, Rodoni BC, et al. Prospects and challenges of implementing DNA metabarcoding for high-throughput insect surveillance. GigaScience [Internet]. 2019;8(8):giz092. Available from: 10.1093/gigascience/giz092

Prince AM, Andrus L. PCR: how to kill unwanted DNA. BioTechniques. 1992;12(3):358–60.

Ramírez-Amaro S, Bassitta M, Picornell A, Ramon C, Terrasa B. Environmental DNA: State-of-the-art of its application for fisheries assessment in marine environments. 2022;9. Available from: https://www.frontiersin.org/articles/10.3389/fmars.2022.1004674;

Revéret A, Rijal DP, Heintzman PD, Brown AG, Stoof-Leichsenring KR, Alsos IG. Environmental DNA of aquatic macrophytes: The potential for reconstructing past and present vegetation and environments. Freshwater Biology [Internet]. 2023 Nov 1;68(11):1929–50. Available from: 10.1111/fwb.14158

Rick TC, Kirch PV, Erlandson JM, Fitzpatrick SM. Archeology, deep history, and the human transformation of island ecosystems. Anthropocene [Internet]. 2013;4:33–45. Available from: http://www.sciencedirect.com/science/article/pii/S2213305413000106

Robbe P, Popitsch N, Knight SJL, Antoniou P, Becq J, He M, et al. Clinical whole-genome sequencing from routine formalin-fixed, paraffin-embedded specimens: pilot study for the 100,000 Genomes Project. Genet Med. 2018;20(10):1196–205.

Roberts P, Hunt C, Arroyo-Kalin M, Evans D, Boivin N. The deep human prehistory of global tropical forests and its relevance for modern conservation. Nature Plants [Internet]. 2017;3(8):17093. Available from: 10.1038/nplants.2017.93

Robinson KM, Prentice C, Clemente-Carvalho R, Hall K, Monteith ZL, Morien E, et al. Paired environmental DNA and dive surveys provide distinct but complementary snapshots of marine biodiversity in a temperate fjord. Environmental DNA [Internet]. 2023 May 1;5(3):597–612. Available from: 10.1002/edn3.423

Rognes T, Flouri T, Nichols B, Quince C, Mahé F. VSEARCH: a versatile open source tool for metagenomics. PeerJ. 2016;4:e2584.

Rohland N, Siedel H, Hofreiter M. Nondestructive DNA extraction method for mitochondrial DNA analyses of museum specimens. BioTechniques [Internet]. 2004;36(5):814–21. Available from: 10.2144/04365ST05

Rojahn J, Gleeson DM, Furlan E, Haeusler T, Bylemans J. Improving the detection of rare native fish species in environmental DNA metabarcoding surveys. Aquatic Conservation: Marine and Freshwater Ecosystems [Internet]. 2021;n/a(n/a). Available from: 10.1002/aqc.3514

Rowe KC, Singhal S, Macmanes MD, Ayroles JF, Morelli TL, Rubidge EM, et al. Museum genomics: lowlJcost and highlJaccuracy genetic data from historical specimens. Mol Ecol Resour [Internet]. 2011 Nov 1;11(6):1082–92. Available from: 10.1111/j.1755-0998.2011.03052.x

Ruppert KM, Kline RJ, Rahman MS. Past, present, and future perspectives of environmental DNA (eDNA) metabarcoding: A systematic review in methods, monitoring, and applications of global eDNA. Global Ecology and Conservation [Internet]. 2019;17:e00547. Available from: https://www.sciencedirect.com/science/article/pii/S2351989418303500

Seersholm FV, Cole TL, Grealy A, Rawlence NJ, Greig K, Knapp M, et al. Subsistence practices, past biodiversity, and anthropogenic impacts revealed by New Zealand-wide ancient DNA survey. Proceedings of the National Academy of Sciences [Internet]. 2018;115(30):7771 LP – 7776. Available from: http://www.pnas.org/content/115/30/7771.abstract

Seersholm FV, Pedersen MW, Søe MJ, Shokry H, Mak SST, Ruter A, et al. DNA evidence of bowhead whale exploitation by Greenlandic Paleo-Inuit 4,000 years ago. Nat Commun [Internet]. 2016 Nov 8;7(1):13389. Available from: 10.1038/ncomms13389

Seymour M, Edwards FK, Cosby BJ, Bista I, Scarlett PM, Brailsford FL, et al. Environmental DNA provides higher resolution assessment of riverine biodiversity and ecosystem function via spatio-temporal nestedness and turnover partitioning. Communications Biology [Internet]. 2021;4(1):512. Available from: 10.1038/s42003-021-02031-2

Shokralla S, Singer GAC, Hajibabaei M. Direct PCR amplification and sequencing of specimens’ DNA from preservative ethanol. BioTechniques [Internet]. 2010;48(3):305–6. Available from: https://www.future-science.com/doi/abs/10.2144/000113362

Spens J, Evans AR, Halfmaerten D, Knudsen SW, Sengupta ME, Mak SST, et al. Comparison of capture and storage methods for aqueous macrobial eDNA using an optimized extraction protocol: advantage of enclosed filter. Methods in Ecology and Evolution [Internet]. 2016;8(5):635–45. Available from: https://besjournals.onlinelibrary.wiley.com/doi/abs/10.1111/2041-210X.12683

Srinivasan M, Sedmak D, Jewell S. Effect of fixatives and tissue processing on the content and integrity of nucleic acids. Am J Pathol. 2002;161:1961–71.

Stadhouders R, Pas SD, Anber J, Voermans J, Mes THM, Schutten M. The effect of primer-template mismatches on the detection and quantification of nucleic acids using the 5’ nuclease assay. The Journal of molecular diagnosticslJ: JMD [Internet]. 2010;12(1):109–17. Available from: https://pubmed.ncbi.nlm.nih.gov/19948821 https://www.ncbi.nlm.nih.gov/pmc/articles/PMC2797725/

Stat M, John J, DiBattista JD, Newman SJ, Bunce M, Harvey ES. Combined use of eDNA metabarcoding and video surveillance for the assessment of fish biodiversity. Conservation Biology [Internet]. 2019;33(1):196–205. Available from: 10.1111/cobi.13183

Stiller M, Sucker A, Griewank K, Aust D, Baretton GB, Schadendorf D, et al. Single-strand DNA library preparation improves sequencing of formalin-fixed and paraffin-embedded (FFPE) cancer DNA. Oncotarget. 2016;7:59115–28.

Stoeckle MY, Mishu MD, Charlop-Powers Z. Improved Environmental DNA Reference Library Detects Overlooked Marine Fishes in New Jersey, United States. Frontiers in Marine Science [Internet]. 2020;7. Available from: https://www.frontiersin.org/articles/10.3389/fmars.2020.00226;

Strickler KM, Fremier AK, Goldberg CS. Quantifying effects of UV-B, temperature, and pH on eDNA degradation in aquatic microcosms. Biological Conservation [Internet]. 2015;183:85–92. Available from: http://www.sciencedirect.com/science/article/pii/S0006320714004637

Suter L, Wotherspoon S, Kawaguchi S, King R, MacDonald AJ, Nester GM, et al. Environmental DNA of Antarctic krill (Euphausia superba): Measuring DNA fragmentation adds a temporal aspect to quantitative surveys. Environmental DNA [Internet]. 2023 Sep 1;5(5):945–59. Available from: 10.1002/edn3.394

Takahashi M, Saccò M, Kestel JH, Nester G, Campbell MA, Heyde M van der, et al. Aquatic environmental DNA: A review of the macro-organismal biomonitoring revolution. Science of the Total Environment [Internet]. 2023 May 15;873:162322. Available from: https://www.sciencedirect.com/science/article/pii/S0048969723009385

Tsuji S, Ushio M, Sakurai S, Minamoto T, Yamanaka H. Water temperature-dependent degradation of environmental DNA and its relation to bacterial abundance. PLOS ONE [Internet]. 2017;12(4):e0176608. Available from: 10.1371/journal.pone.0176608

Vasiliadis M, Freer JJ, Collins MA, Cleary AC. Assessing the trophic ecology of Southern Ocean Myctophidae: the added value of DNA metabarcoding. Canadian Journal of Fisheries and Aquatic Sciences [Internet]. 2024 Feb 1;81(2):166–77. Available from: 10.1139/cjfas-2023-0079

Wang Z, Liu X, Liang D, Wang Q, Zhang L, Zhang P. VertU: universal multilocus primer sets for eDNA metabarcoding of vertebrate diversity, evaluated by both artificial and natural cases. Frontiers in Ecology and Evolution [Internet]. 2023;11. Available from: https://www.frontiersin.org/articles/10.3389/fevo.2023.1164206;

Williams C, Pontén F, Moberg C, Söderkvist P, Uhlén M, Pontén J, et al. A high frequency of sequence alterations is due to formalin fixation of archival specimens. Am J Pathol. 1999;155:1467–71.

Willis KJ, Araújo MB, Bennett KD, Figueroa-Rangel B, Froyd CA, Myers N. How can a knowledge of the past help to conserve the future? Biodiversity conservation and the relevance of long-term ecological studies. Philosophical transactions of the Royal Society of London Series B, Biological sciences [Internet]. 2007;362(1478):175–86. Available from: https://www.ncbi.nlm.nih.gov/pubmed/17255027 https://www.ncbi.nlm.nih.gov/pmc/articles/PMC2311423/

Wood JR, Scofield RP, Hamel J, Lalas C, Wilmshurst JM. Short communication: Bone stable isotopes indicate a high trophic position for new zealand’s extinct south island adzebill (Aptornis defossor) (Gruiformes: Aptornithidae). New Zealand Journal of Ecology. 2017;41(2):240–4.

Woods BL, Putte APV de, Hindell MA, Raymond B, Saunders RA, Walters A, et al. Species distribution models describe spatial variability in mesopelagic fish abundance in the Southern Ocean. Frontiers in Marine Science [Internet]. 2023;9. Available from: https://www.frontiersin.org/articles/10.3389/fmars.2022.981434;

Wright ES. Using DECIPHER v2. 0 to analyze big biological sequence data in R. The R Journal. 2016;8(1).

Wulff J. Sponge Contributions to the Geology and Biology of Reefs: Past, Present, and Future. In Dordrecht: Springer Netherlands; 2016. p. 103–26. Available from: 10.1007/978-94-017-7567-0_5

Zimmermann J, Hajibabaei M, Blackburn DC, Hanken J, Cantin E, Posfai J, et al. DNA damage in preserved specimens and tissue samples: a molecular assessment. Front Zoo[l [Internet]. 2008 Oct 23;5(1):18. Available from: 10.1186/1742-9994-5-18

