## Supplement 1 for "Unlocking Antarctic molecular time-capsules – recovering historical environmental DNA from museum-preserved sponges"

### Supplement 1: NIWA Invertebrate Collection sponge specimen metadata

Supplementary Table 6: NIWA Invertebrate Collection sponge specimen metadata, including the NIWA catalogue number (Cat #), preservation technique used to store the specimen (Preservation), taxonomic information of order (Order) and species (Species) for each specimen, collection year (Year), sampling location (Latitude, Longitude), depth of the sponge upon collection (Depth), sample collection method (Gear), and specimen ID used through the manuscript (ID). Latitude and longitude coordinates of specimens provided by CCAMLR fisheries are truncated to the nearest degree according to the conditions for the provision of this data in publications and protect commercial interest.

| Cat # | Preservation | Order | Species | Year | Latitude | Longitude | Depth | Gear | ID |
| --- | --- | --- | --- | --- | --- | --- | --- | --- | --- |
| 36822 | Dry | Poecilosclerida | *Inflatella belli* | 2008 | -76.1930 | 176.2961 | 447 | Trawl | PMD1 |
| 36826 | Dry | Lyssacinosida | *Rossella nuda* | 2008 | -76.1930 | 176.2961 | 447 | Trawl | PMD2 |
| 35650 | Dry | Lyssacinosida | *Rossella villosa* | 2008 | -74.5817 | 170.2498 | 285 | Trawl | PMD3 |
| 35590 | Dry | Lyssacinosida | *Rossella villosa* | 2008 | -74.1111 | 170.7960 | 639 | Trawl | PMD6 |
| 95376 | Dry | Tetractinellida | *Cinachyra barbata* | 1960 | -74.0550 | 179.3500 | 267 | Trawl | PMD4 |
| 37384 | Dry | Suberitida | *Homaxinella sp.* | 2008 | -72.0926 | 175.5716 | 1,567 | Trawl | PMD7 |
| 37655 | Dry | Suberitida | *Homaxinella sp.* | 2008 | -72.0755 | 172.9043 | 535 | Trawl | PMD5 |
| 37237 | Dry | Suberitida | *Homaxinella sp.* | 2008 | -72.3395 | 175.5322 | 945 | Trawl | PMD8 |
| 37457 | Dry | Suberitida | *Homaxinella sp.* | 2008 | -72.9385 | 173.3023 | 1,431 | Trawl | PMD9 |
| 37464 | Dry | Suberitida | *Homaxinella sp.* | 2008 | -72.9385 | 173.3023 | 1,431 | Trawl | PMD10 |
| 145261 | Frozen | Lyssacinosida | *Rossella nuda* | 2009 | -71 | 171 | 1,120 | Longline | PMF1 |
| 145244 | Frozen | Lyssacinosida | *Rossella villosa* | 2009 | -72 | 172 | 1,307 | Longline | PMF2 |
| 146336 | Frozen | Suberitida | *Homaxinella sp.* | 2010 | -75 | -177 | 642 | Longline | PMF7 |
| 146346 | Frozen | Suberitida | *Homaxinella sp.* | 2010 | -75 | -177 | 813 | Longline | PMF8 |
| 146328 | Frozen | Suberitida | *Homaxinella sp.* | 2010 | -75 | 176 | 760 | Longline | PMF9 |
| 146332 | Frozen | Suberitida | *Homaxinella sp.* | 2010 | -75 | 177 | 707 | Longline | PMF3 |
| 146334 | Frozen | Suberitida | *Homaxinella sp.* | 2010 | -75 | -177 | 1,227 | Longline | PMF10 |
| 131631 | Frozen | Poecilosclerida | *Inflatella belli* | 2009 | -72 | 176 | 1,265 | Longline | PMF4 |
| 146356 | Frozen | Lyssacinosida | *Rossella villosa* | 2010 | -75 | -177 | 1,227 | Longline | PMF6 |
| 146319 | Frozen | Tetractinellida | *Cinachyra barbata* | 2011 | -73 | -177 | 1,162 | Longline | PMF5 |
| 29076 | Ethanol | Lyssacinosida | *Rossella villosa* | 2004 | -72.3209 | 170.4751 | 272 | Trawl | PME1 |
| 29113 | Ethanol | Lyssacinosida | *Rossella villosa* | 2004 | -72.3418 | 170.4429 | 203 | Trawl | PME6 |
| 13613 | Ethanol | Lyssacinosida | *Rossella nuda* | 2008 | -72.5903 | 175.3423 | 479 | Trawl | PME2 |
| 36471 | Ethanol | Tetractinellida | *Cinachyra barbata* | 2008 | -76.8330 | 179.9500 | 664 | Trawl | PME3 |
| 35450 | Ethanol | Suberitida | *Homaxinella sp.* | 2008 | -73.1245 | 174.3205 | 321 | Trawl | PME7 |
| 35613 | Ethanol | Suberitida | *Homaxinella sp.* | 2008 | -74.1111 | 170.7960 | 639 | Trawl | PME4 |
| 37229 | Ethanol | Suberitida | *Homaxinella sp.* | 2008 | -72.5850 | 175.3083 | 467 | Trawl | PME8 |
| 50605 | Ethanol | Suberitida | *Homaxinella sp.* | 2008 | -72 | 172 | 1,050 | Longline | PME9 |
| 37362 | Ethanol | Suberitida | *Homaxinella sp.* | 2009 | -72.0927 | 175.5717 | 1,567 | Trawl | PME10 |
| 35705 | Ethanol | Poecilosclerida | *Inflatella belli* | 2008 | -74.5816 | 170.2498 | 285 | Trawl | PME5 |

### Supplement 2: Historical eDNA extraction protocol from museum-stored sponges

From each of the 30 sponge specimens, five tissue biopsies (approximate size of tissue biopsy: 0.5 cm^3^) were collected at the NIWA Invertebrate Collection in Wellington, New Zealand. All tissue biopsies were transported to the eDNA laboratory facilities at the Portobello Marine Laboratory, University of Otago, New Zealand. Transport conditions matched the initial storage condition of each specimen, i.e., frozen specimens were transported and kept frozen, dried specimens were transported dried, and ethanol submerged specimens were transported in ethanol. DNA was extracted from all five tissue biopsies of each of the 30 sponge specimens using the Qiagen DNeasy Blood & Tissue Kit (Qiagen GmbH, Germany; Cat# 69506) principally following the manufacturer’s protocol, but with slight modifications. Each extraction round included 23 tissue biopsies and one negative extraction control. The exact DNA extraction protocol is detailed below:

1. From each sponge specimen, biopsy 5 tissue samples with a size of approximately 0.5 cm^3^.
2. Place each tissue biopsy in a separate 2 ml Eppendorf tube.
3. Add 720 µl Buffer ATL and 80 µl Proteinase K.
4. Vortex at maximum speed for 1 minute to mix sample.
5. Pulse spin Eppendorf tubes using benchtop centrifuge to move all liquid to the bottom.
6. Incubate samples at 56°C overnight.
7. Remove samples from incubator.
8. Pipette mix 10 times to homogenize sample.
9. Place Buffer AE in incubator at 56°C.
10. Centrifuge samples at 6,000 x g for 1 minute at room temperature.
11. Pipette 600 µl of supernatant (be careful not to include spicules) into a new 2 ml Eppendorf tube.
12. Add 600 µl Buffer AL.
13. Vortex at maximum speed for 15 seconds.
14. Add 600 µl 100% Ethanol (best kept frozen prior to DNA extraction).
15. Vortex samples at maximum speed for 15 seconds.
16. Pipette the mixture into a DNeasy Mini Spin Column (maximum 620 µl at a time).
17. Centrifuge samples at 6,000 x g for 1 minute.
18. Discard the flow-through and collection tube.
19. Place the DNeasy Mini Spin Column in a new 2 ml collection tube.
20. Repeat steps 16 – 19 until all the mixture has gone through the DNeasy Mini Spin Column, changing the collection tube every time.
21. Add 500 µl Buffer AW1.
22. Centrifuge samples for 1 minute at 6,000 x g.
23. Discard the flow-through and collection tube.
24. Place the DNeasy Mini Spin Column in a new 2 ml collection tube.
25. Add 500 µl Buffer AW2.
26. Centrifuge samples for 3 minutes at 20,000 x g.
27. Discard the flow-through and collection tube.
28. Place the DNeasy Mini Spin Column in a new 2 ml collection tube.
29. Centrifuge samples for 1 minute at 20,000 x g to remove any access liquid.
30. Discard the flow-through and collection tube.
31. Transfer the DNeasy Mini Spin Column in a 2 ml Eppendorf tube with caps removed.
32. Add 100 µl Buffer AE to the center of the membrane (preheat Buffer AE to 56°C; see step 6)
33. Incubate samples for 1 minute at room temperature.
34. Centrifuge samples for 1 minute at 6,000 x g.
35. Repeat 100 µl elution step to a total elution volume of 200 µl.
36. Transfer eluate to a new 1.5 ml Eppendorf tube.
37. Store DNA at -20°C until further processing.

### Supplement 3: IDTAXA reference database for taxonomy assignment

Please find the code on how to create the IDTAXA reference database on the GitHub repository <https://github.com/gjeunen/marsden_obj1_preservationMethod>. The reference database used to assign a taxonomic ID to each ZOTU sequence in this dataset can be found as an additional supplemental file to this document labeled “supplement_3_idtaxa.fasta”.

### Supplement 4: ZOTU frequency table

The ZOTU frequency table can be generated from the raw sequence files and bioinformatic processing described in the GitHub repository (<https://github.com/gjeunen/marsden_obj1_preservationMethod>). Additionally, the ZOTU frequency table used in this manuscript can be found as an extra supplemental file to this document labeled “supplement_4_frequency_table.xlsx”.

### Supplement 5: Rarefaction curves

No significant correlation was observed between sequencing depth and number of detected taxa according to a Pearson correlation test (t = 0.805, df = 132, p-value = 0.422; Supplementary Figure 5.1). Rarefaction curves were generated from the unfiltered frequency table to assess sequencing coverage using the *vegan v* 2.5-7 and ampvis2 R packages. Overall, samples achieved sufficient sequencing coverage based on the plateauing of rarefaction curves (Supplementary Figure 5.2) and mean number of reads per sample ± *SD*: 73,357 ± 25,183.

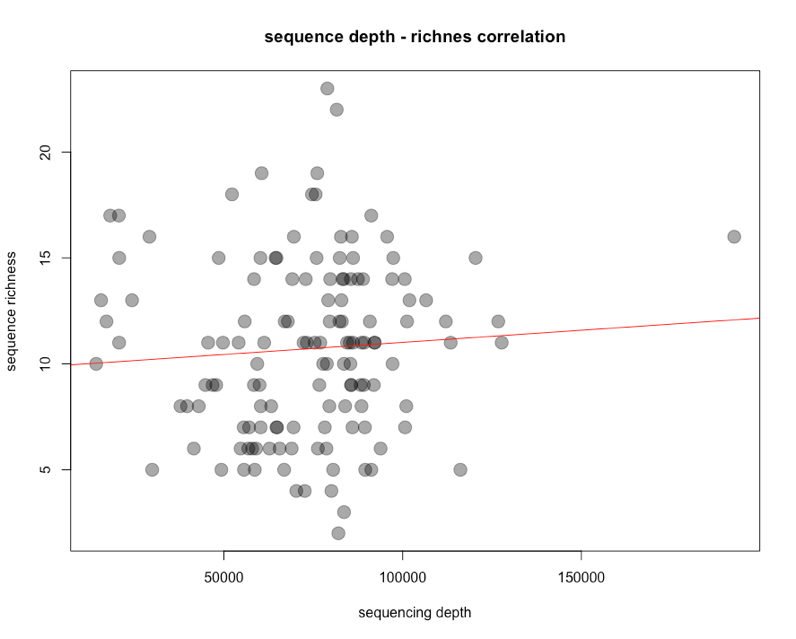

Supplementary Figure 5.1: Correlation analysis plot displaying the number of sequences on the x-axis and number of detected taxa on the y-axis for each sample. Regression line is indicated in red, while samples are represented as black circles.

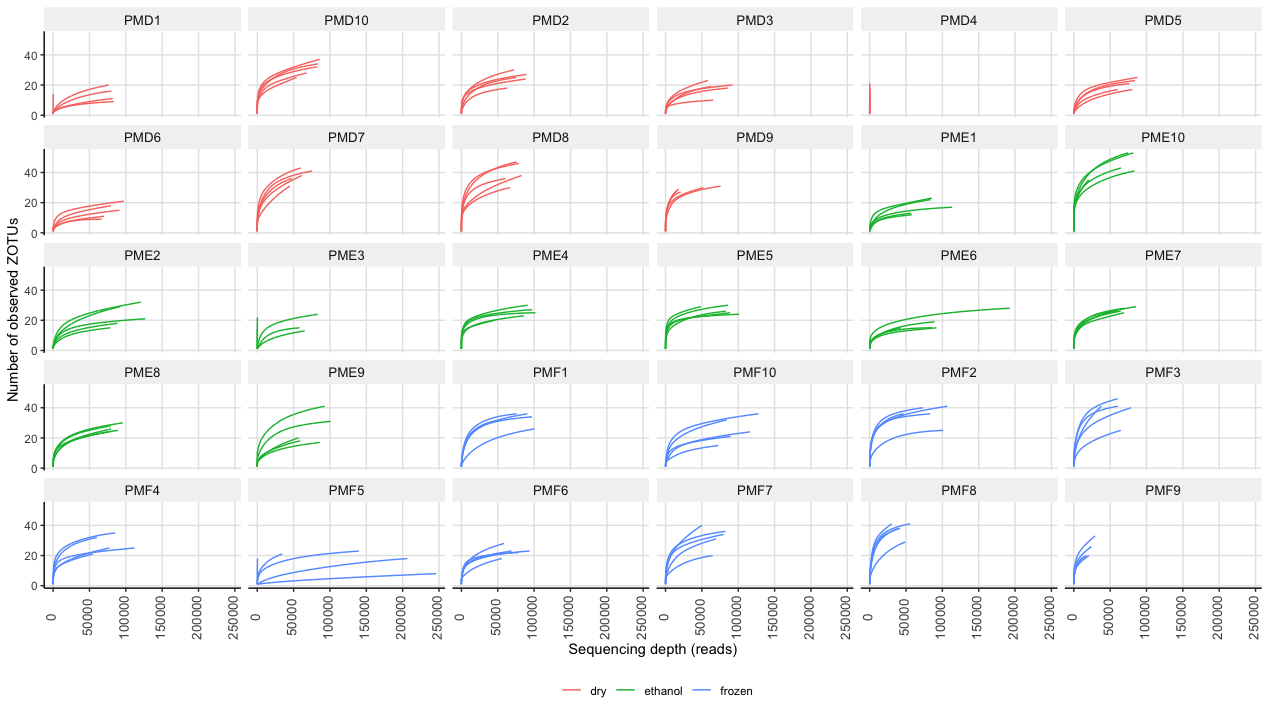

Supplementary Figure 5.2: Rarefaction plots facetted by sponge specimen, with sequencing depth on the x-axis and number of detected ZOTUs on the y-axis. Dried sponges are coloured red, ethanol-submerged sponges are coloured green, and frozen sponges are coloured blue. Specimen labels follows formatting in Supplement 1.

### Supplement 6: Species accumulation curves

Due to the multiple biopsies per sponge, transforming the count table to presence-absence data enables the dataset to be secondarily transformed to an incidence frequency table when merging data from different biopsies per sponge, resulting in a semi-quantitative dataset whereby values range from 0 (ZOTU not detected within a single biopsy) to 5 (ZOTU detected in all biopsies). Through inter- and extrapolation calculations using the iNEXT.3D R package, species accumulation curves can be drawn (Supplemental Figure 6). The plateauing of species accumulation curves revealed five biopsies to be sufficient to recover most of the fish diversity held within a sponge.

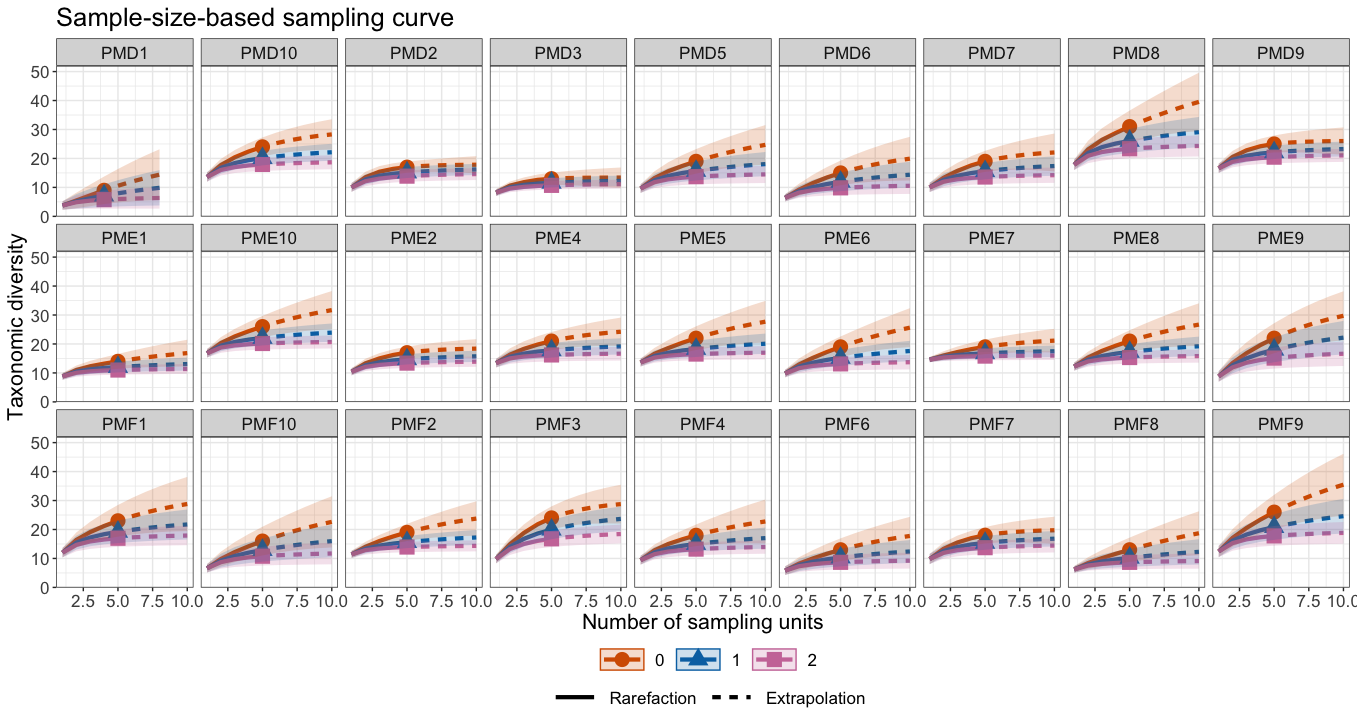

Supplementary Figure 6: Species accumulation curves facetted by sponge specimens for three orders of Hill numbers, including q = 0 (species richness; orange), q = 1 (exponential of Shannon entropy; blue), and q = 2 (inverse of Simpson concentration; purple). Specimen labels follows formatting in Supplement 1.

### Supplement 7: Estimated replication analysis without removal of outlier

The estimated replication, i.e., number of biopsies per sponge, needed to recover 90% of the fish diversity, based on inter- and extrapolation calculations, did not differ significantly between preservation methods according to a Welch’s ANOVA (*F*_2,15_ = 2.20, *p* = 0.144; Supplemental Figure 7).

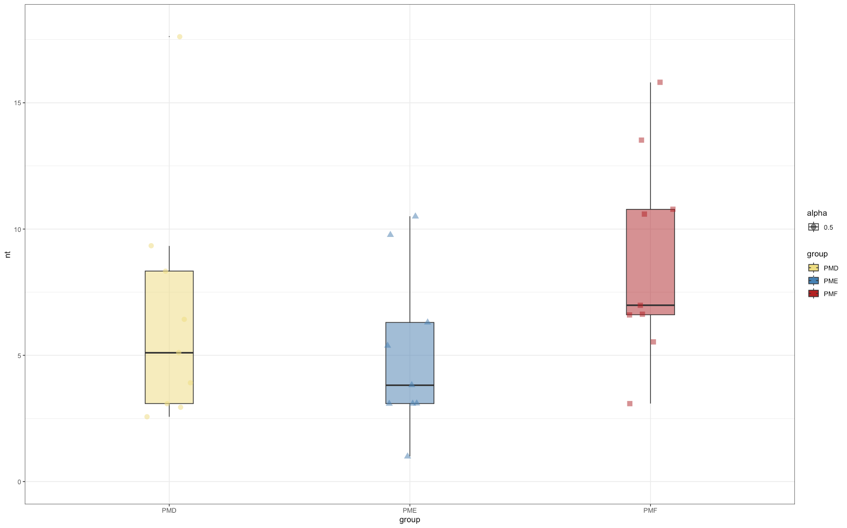

Supplementary Figure 7: Boxplots depicting the estimated tissue biopsies required to recover 90% of the fish diversity between the three preservation methods, including dry (yellow), ethanol (blue), and frozen (red). The median is indicated by a black line within each boxplot. Samples are indicated by coloured dots, including circle (dry), triangle (ethanol), and square (frozen).

### Supplement 8: *In silico* PCR analysis of myctophids

The *in silico* PCR analysis was conducted in CRABS *v* 0.1.5 by downloading 444 Myctophidae 16S rRNA sequences. The sequences were analyzed using the standard protocol described on the GitHub repository <https://github.com/gjeunen/reference_database_creator>, whereby the primers used in this study were selected as the primer set for the *in silico* PCR analysis (Fish16SF: 5’-GACCCTATGGAGCTTTAGAC-3'; Fish16S2R: 5’-CGCTGTTATCCCTADRGTAACT-3'). Supplemental Figure 8 was generated using the ‘*visualisation*’ function within CRABS.

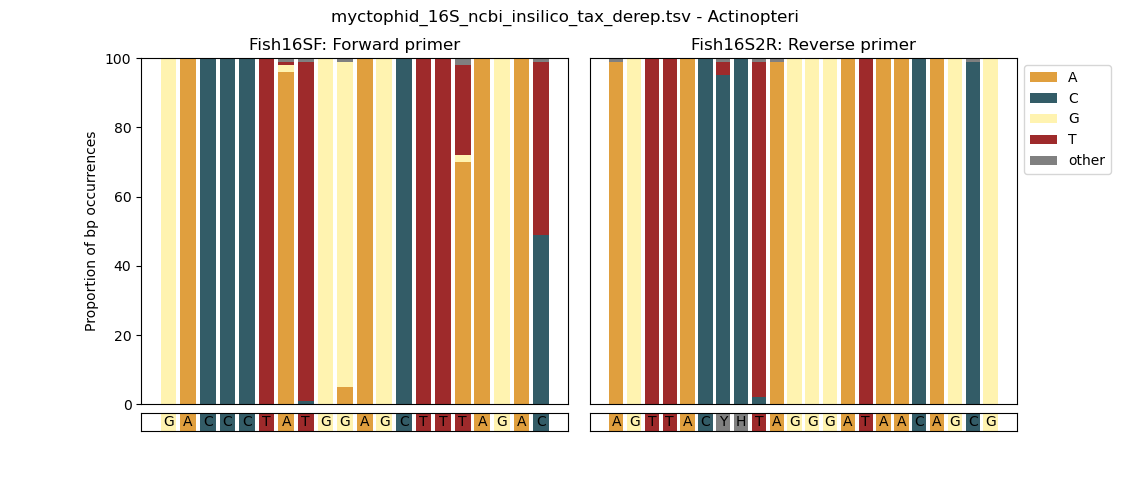

Supplementary Figure 8: In silico PCR analysis of 444 Myctophidae sequences for the 16S rRNA gene. Forward and reverse primer sequences are displayed on the bottom, while proportion of basepair occurrences are displayed on top. Colour coding of base pairs identifies two sites within the 3’ end of the forward primer as high proportion mismatch sites for Myctophidae.
